# *In situ* characterization of calcium fluxes in astrocytic mitochondria from the mouse striatum and hippocampus

**DOI:** 10.1101/2020.06.21.163618

**Authors:** Taylor E. Huntington, Rahul Srinivasan

## Abstract

Astrocytes govern critical aspects of brain function via Ca^2+^ signals, the majority of which associate with mitochondria. However, little is known with regard to *in situ* sources, kinetics or mechanisms of Ca^2+^ influx in astrocytic mitochondria. To address this knowledge gap, we expressed the genetically encoded calcium indicator, GCaMP6f within the mitochondrial matrix of adult mouse astrocytes in the dorsolateral striatum (DLS) and hippocampus (HPC). We found spontaneous Ca^2+^ events in astrocytic mitochondria with subcellular differences between somatic, branch, and branchlet mitochondria, as well as inter-regional differences between astrocytes in the DLS and HPC. We also found a strong dependency of spontaneous mitochondrial Ca^2+^ fluxes on endoplasmic reticulum stores, the surprising lack of a major role for the mitochondrial calcium uniporter, MCU, and dual mitochondrial Ca^2+^ responses with multiple neurotransmitter agonists. Together, our findings provide a foundational understanding of mechanisms for Ca^2+^ influx in astrocytic mitochondria within disease-relevant brain regions.

## Introduction

Once regarded as mere supporting cells, astrocytes have recently emerged as critical players in governing multiple aspects of brain physiology. Among their many functions, these cells modulate neural activity^1,2^, control synapse formation^3^, maintain K^+^ homeostasis and neuronal excitability^4,5^, regulate neurovasculature^6,7^, and provide ~20% of the total energy required by the brain^8,9^.

The multi-faceted effects of astrocytes on nervous system function are thought to depend on spontaneous Ca^2+^ signals in astrocytic somata and the fine processes within their territories^10–14^. Strong evidence for a central role played by astrocytic Ca^2+^ signals in regulating neural function comes from multiple reports showing that astrocytic Ca^2+^ signals are enhanced by behavioral stimuli such as forced locomotion^15^, or an air puff-induced startle response^16^, and are robustly modulated by drugs affecting the central nervous system, such as anesthetics^17,18^ and amphetamine^19^. Importantly, the disruption of Ca^2+^ fluxes specifically in astrocytes causes repetitive behaviors in mice^20^, potentiates short-term plasticity in the hippocampus^21^, and inhibits metabolic coupling between astrocytes and neurons^22^. Together these studies show that astrocytic Ca^2+^ signals are capable of influencing critical aspects of brain activity and consequently, behavioral outputs during health and disease. Therefore, developing an understanding of the mechanisms underlying astrocytic Ca^2+^ signals is important from a basic, as well as a translational perspective.

A recent study showed that the vast majority of spontaneous Ca^2+^ events within astrocytic processes *in vivo* occur due to brief periods of Ca^2+^ efflux through the mitochondrial permeability transition pore (mPTP), and are abnormally increased in a mouse model of amyotrophic lateral sclerosis ^23^. This suggests that mitochondrial Ca^2+^ fluxes in astrocytes are not only critical for normal brain function, but can also significantly influence neurodegenerative processes. The central role played by astrocytic mitochondria during neurodegeneration is further highlighted by independent studies showing that disruption of astrocytic mitochondria exacerbates neurodegeneration in cerebellar Purkinje neurons^24^, attenuates neuroprotection following ischemia^25^, and even slows the recovery of mice from anesthesia^26^. Despite the many important roles ascribed to astrocytic mitochondria, and specifically to mitochondrial Ca^2+^ fluxes in astrocytes during health and disease, we know very little with regard to the subcellular characteristics of astrocytic mitochondria *in situ*, whether or not Ca^2+^ spontaneously fluxes into the mitochondria of astrocytes, and the extent to which astrocytic mitochondrial Ca^2+^ events respond to neurotransmitter agonists.

To address these broadly significant questions, we generated an adeno-associated virus (AAV)-based genetically encoded calcium indicator (GCaMP6f) specifically targeted to the matrix of astrocytic mitochondria and directly measured mitochondrial Ca^2+^ influx in striatal and hippocampal astrocytes *in situ* from adult mouse brain slices. Our experiments reveal that astrocytic mitochondria in the dorsolateral striatum (DLS) and hippocampus (HPC) display unique characteristics with regard to their morphological features, the kinetics of spontaneous mitochondrial Ca^2+^ events, sources and portals for entry of Ca^2+^ into mitochondria, and the response of astrocytic mitochondrial Ca^2+^ events to neurotransmitter agonists. These findings have important implications for developing a basic understanding of how mitochondrial Ca^2+^ events in astrocytes can influence normal brain physiology, and thereby contribute to a number of neuropathological processes, including neurodegeneration.

## Materials and Methods

### Generation of AAV 2/5 GfaABC_1_D-mito-7-GCaMP6f

The GfaABC_1_D-mito-7-GCaMP6f construct was generated by Vector Builder (Chicago, IL) using the Gateway cloning method^27^. Entry clones containing the mito-7 signaling sequence (87 bp), derived from COX8A and the GfaABC1D promoter (877 bp) were generated and recombined into a destination vector to create the construct. The GfaABC1D-mito-7-GCaMP6f cassette was then introduced into a pZac2.1 plasmid to create pZac2.1 GfaABC1D-mito-7-GCaMP6f, and this plasmid was used to generate the AAV 2/5 GfaABC_1_D-mito-7-GCaMP6f viral vector.

### Mice

Male and female C57BL/6 WT breeders were obtained from Taconic and bred in house. Mice used for MCU^−/−^ experiments were of a CD1 background and purchased from Texas A&M Institute for Genomic Medicine (TIGM). MCU^−/−^ mice were generated via gene trap method by integrating a retroviral trapping vector into the first intron of the CCDC109A locus^28^. WT CD1 littermates were used as controls for MCU^−/−^ experiments.

Mice were housed on a 12 h light/dark cycle with *ad libitum* access to food and water. For Ca^2+^ imaging, mice were injected with AAV2/5 GfaABC_1_D-mito-7-GCaMP6f at approximately 8 weeks old and imaged 3 weeks later. All animal experiments were conducted in accordance with Texas A&M University IACUC guidelines.

### Immunostaining

For staining mouse DLS and HPC sections, mice were transcardially perfused with 10% formalin and brains were extracted. Brains were postfixed in 10% formalin overnight at 4°C and cryoprotected in 30% sucrose for 48-72 hrs. 40 μm thick coronal sections were obtained using a cryostat microtome (Leica) and preserved in 0.01% sodium azide + PBS at 4°C until use. Immunohistochemistry was performed using previously published techniques^16,29^. Briefly, sections were washed 3x for 10 min in 1X TBS, then blocked for 1 hr at RT in 1X TBS solution containing 5% NGS + 0.25% Triton-X-100. Sections were incubated overnight at 4°C in primary antibodies diluted in blocking solution. The following primary antibodies were used: chicken anti-GFP (1:2000; Abcam ab13970) and mouse anti-pyruvate dehydrogenase (PDH) (1:1000, Abcam ab110333). The following day sections were washed 3x for 10 min each in 1X TBS and incubated with appropriate secondary antibodies in blocking solution for 2 hr at RT. The following secondary antibodies were used: Alexa-488 goat anti-chicken (1:2000; Abcam ab150176) and Alexa-594 goat anti-mouse (1:2000; Abcam ab150120). Sections were rinsed 3x for 10 min in 1X TBS and then mounted on microscope slides in Fluoromount (Diagnostic Biosystems; K024) for imaging.

### Stereotaxic surgery

Surgical procedures were conducted as previously described^16^. Briefly, surgeries were performed under general anesthesia using continuous isoflurane (induction at 5%, maintenance at 1-2% vol/vol) administered via a syringe injection system (Kent Scientific). Following head fixation on a stereotaxic frame (David Kopf Instruments, Tujunga CA), a 2-3 mm diameter craniotomy was performed using a high-speed dental drill (Foredom) and 0.9% saline was intermittently applied to reduce heating caused by the drill. Glass injection pipettes (World Precision Instruments, 1B100-4) were pulled using a Sutter P-2000 laser puller and beveled using a Narishige EG-45 grinder. 2 × 10^9^ genome copies (gc) of AAV2/5 GfaABC_1_D-mito7-GCaMP6f was injected in a 2 μL volume using a syringe pump (Harvard Apparatus). The injection pipette was left in place for ~10 min after injection and then gradually withdrawn. Surgical wounds were closed with non-absorbable 5-0 sutures (Ethicon, 682G) and mice were sacrificed 20-23 days post-surgery for Ca^2+^ imaging. The injection coordinates for the DLS were as follows: anterior to bregma, +0.9 mm; lateral to bregma, +1.8 mm; and ventral to pial surface, +2.5 mm. Coordinates for the CA1 region of HPC were as follows: posterior to bregma, −2.0 mm; lateral to bregma, +1.5 mm; and ventral to pial surface, −1.6 mm.

### Ca^2+^ imaging in acute mouse brain slices

300 μm thick coronal slices of either the DLS or HPC were cut in a slicing solution consisting of (in mM): 194 sucrose, 30 NaCl, 4.5 KCl, 1.2 NaH_2_PO_4_, 26 NaHCO_3_, 10 D-glucose, and 1 MgCl_2_ and saturated with 95% O_2_ and 5% CO_2_ (pH 7.2). Incubation and recording were performed in artificial cerebrospinal fluid (ACSF) comprising (in mM): 126 NaCl, 2.5 KCl, 1.24 NaH_2_PO_4_, 26 NaHCO_3_, 10 D-glucose, 2.4 CaCl_2_, and 1.3 MgCl_2_ saturated with 95% O_2_ and 5% CO_2_ (pH 7.4). Slices were incubated in ACSF at 34°C for 10 min and then incubated in ACSF with 150 nM of MitoTracker Deep Red (MTDR) (Invitrogen) at 34°C for 1 hr, just prior to imaging. 150 nM MTDR^30,31^ was used as an optimal concentration in acute slices and was determined as an appropriate concentration based on preliminary testing with concentrations ranging from 50 nM-300 nM.

Imaging was performed as previously described^16^. Briefly, slices were imaged using an Olympus FV1200 upright laser-scanning confocal microscope with a 40X water immersion objective lens, numerical aperture (NA) of 0.8, 488 nm, and 633 nm laser lines. For each imaging session, the 488 nm line intensity was set at 10% maximum output to visualize GCaMP6f fluorescence, and the 633 nm laser line at 1-5% maximum output to visualize MTDR fluorescence. Confocal parameters (high voltage, gain, offset, laser power, and aperture diameter) were maintained constant for all GCaMP6f imaging and were optimized for MTDR during each imaging session. Ca^2+^ events were recorded at either 800 msec/frame (Figs. 1, 3-10) or 1 frame/sec (FPS) (Fig. 2).

**Figure 1.**
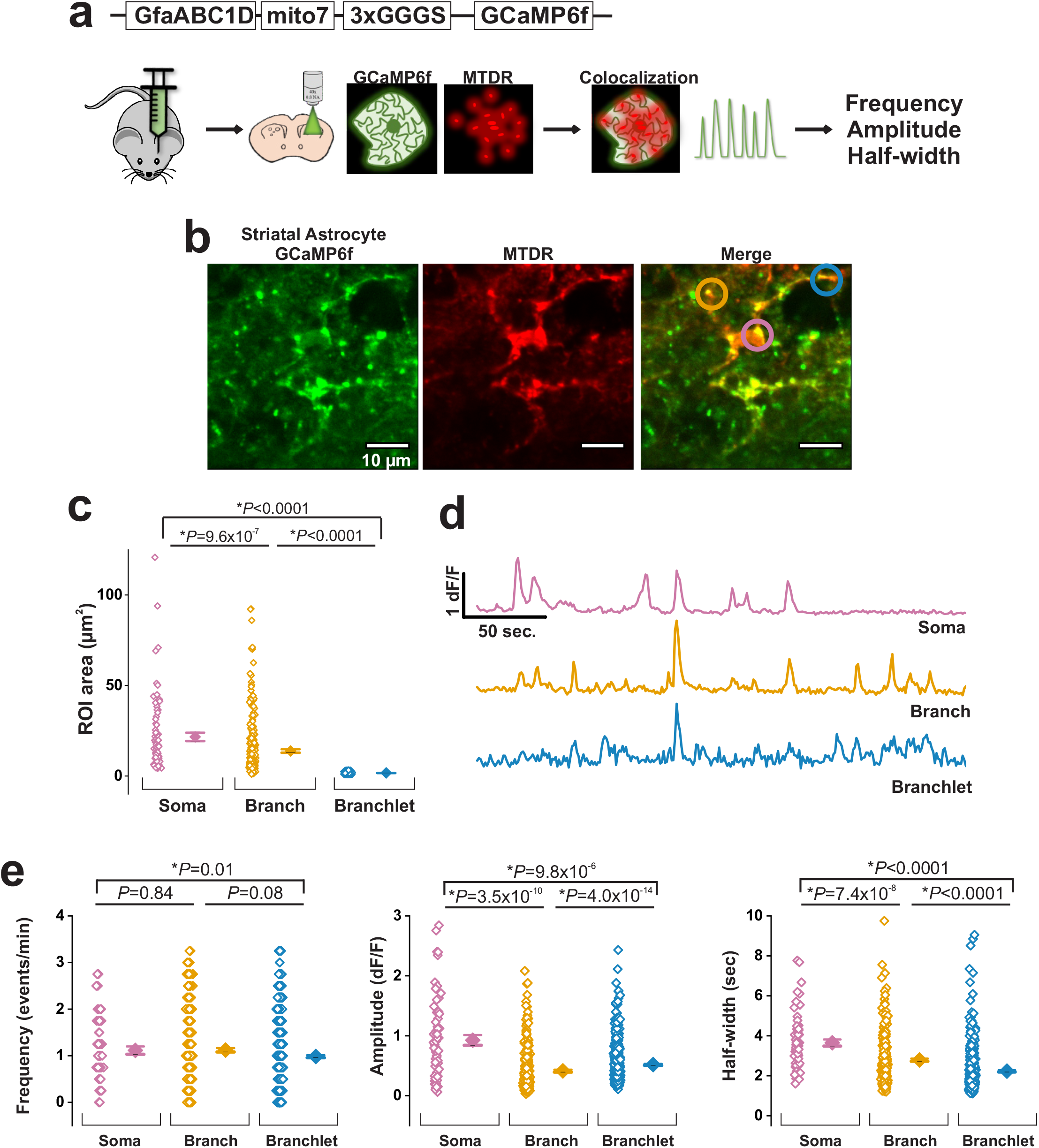
Spontaneous Ca^2+^ events in striatal astrocytic mitochondria visualized with AAV 2/5-GfaABC_1_D-mito-7-GCaMP6f. **a**, Schematic of the GfaABC_1_D-mito-7-GCaMP6f construct and experimental approach. **b**, Representative time compressed confocal images of a striatal astrocyte expressing GCaMP6f (green) and MitoTracker Deep Red (MTDR, red) from live brain slices of WT C57BL/6 mice. Merged image shows somatic, branch, and branchlet mitochondrial regions of interest (ROIs) in magenta, orange, and blue circles respectively. Scale bar = 10 μm. **c**, Areas of somatic, branch and branchlet mitochondria in DLS astrocytes (56 astrocytes and 15 mice; n= 67 somatic, 336 branch, and 605 branchlet mitochondria). **d**, Representative traces of spontaneous mitochondrial Ca^2+^ events in somatic (magenta), branch (orange) and branchlet (blue) mitochondria. **e**, Population data and mean values for spontaneous Ca^2+^ event frequency (left), amplitude (middle), and half-width (right) in striatal astrocytic mitochondria from 56 astrocytes and 15 mice (n= 67 somatic, 336 branch, and 605 branchlet mitochondria). Errors are ± s.e.m. p-values are based on Mann-Whitney test.

**Figure 2.**
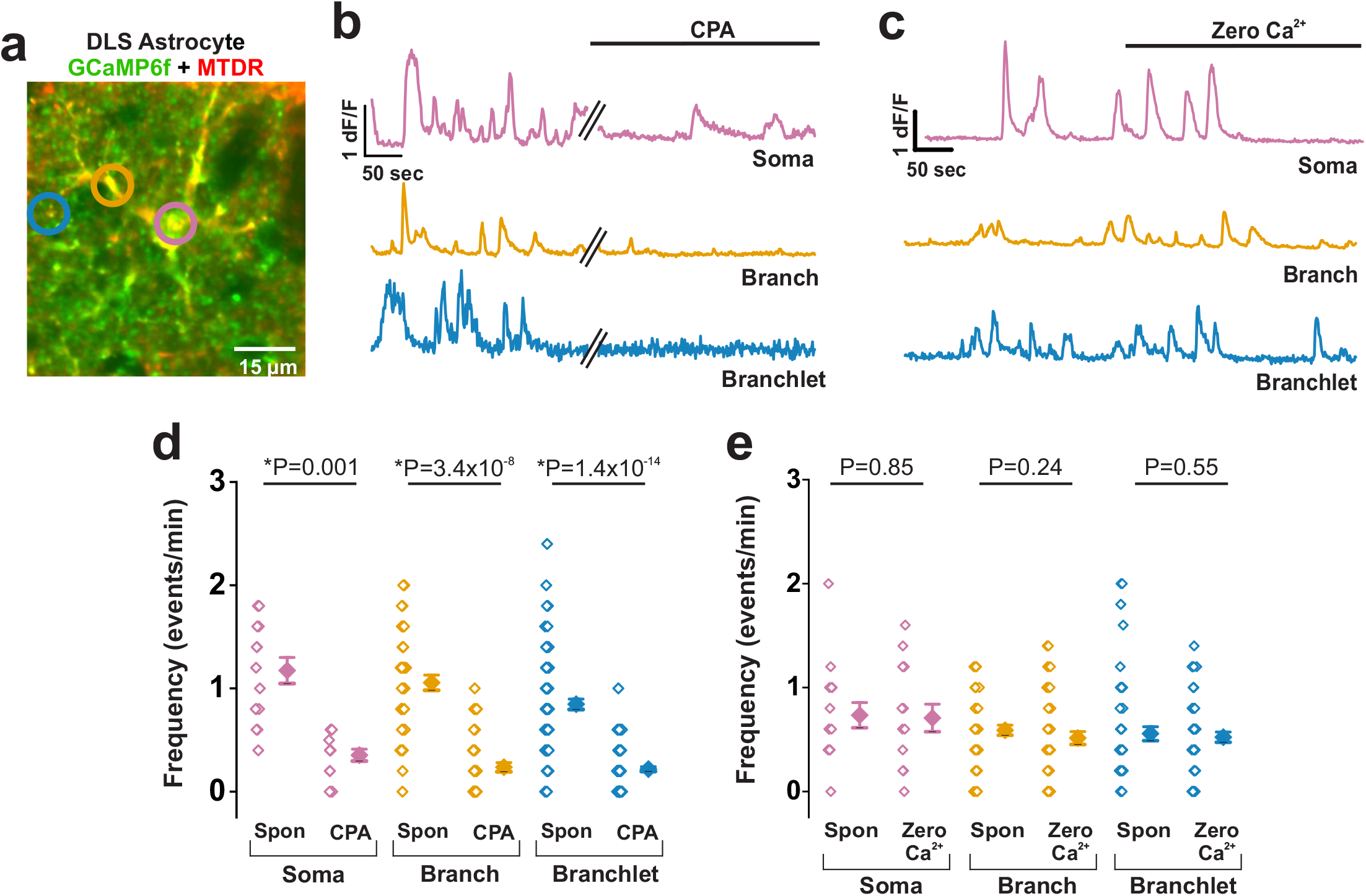
Spontaneous mitochondrial Ca^2+^ events in DLS astrocytes depend on ER-localized Ca^2+^ stores. **a**, Representative merged time compressed confocal image of a DLS astrocyte expressing GCaMP6f (green) and MTDR (red). Somatic, branch, and branchlet ROIs are shown by magenta, orange, and blue circles respectively. Scale bar = 15 μm. **b**, Representative Ca^2+^ event traces of mitochondria in each subpopulation in response to CPA. **c**, Representative Ca^2+^ event traces from DLS astrocytic mitochondria before and after Zero Ca^2+^. **d**, Population data and mean values showing changes in Ca^2+^ event frequency in mitochondria before and after CPA (7 astrocytes and 4 mice; n= 15 somatic, 43 branch, and 86 branchlet mitochondria). **e**, Ca^2+^ event frequency in mitochondria from each subpopulation before and after Zero Ca^2+^ (8 astrocytes and 4 mice; n= 12 somatic, 45 branch, and 59 branchlet mitochondria). Errors are ± s.e.m. p-values are based on Wilcoxon Signed Rank test for somatic mitochondria before and after CPA administration. All other data sets (CPA and Zero Ca^2+^) were subject to paired t-test.

**Figure 3.**
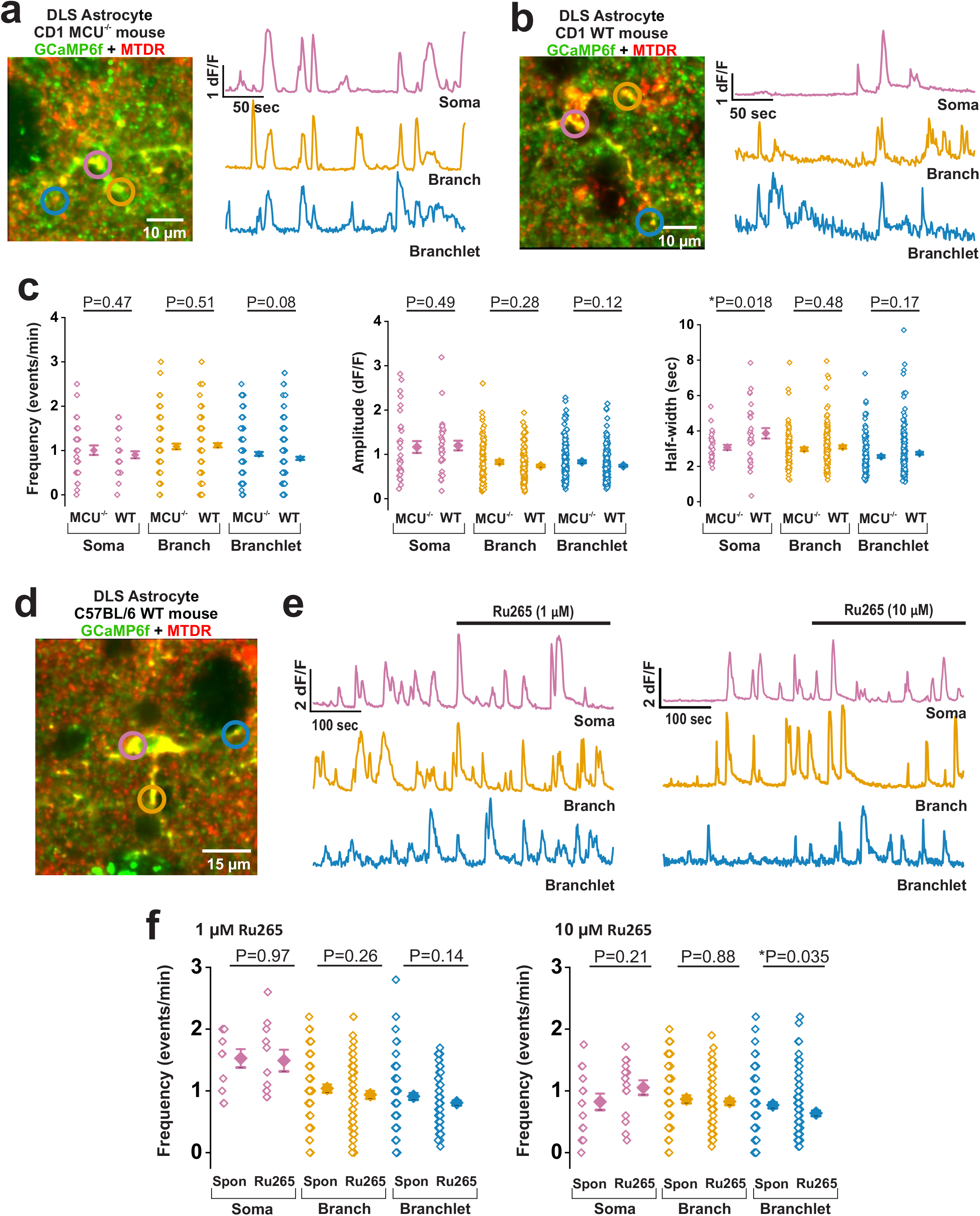
Ca^2+^ fluxes in astrocytic mitochondria do not depend on MCU. Representative merged time compressed confocal images of GCaMP6f + MTDR labeled DLS astrocytes and traces of spontaneous astrocytic mitochondrial Ca^2+^ events from **a**, CD1 MCU^−/−^ and **b**, CD1 WT littermates. In each case, ROIs and corresponding traces for somatic (magenta), branch (orange), and branchlet (blue) mitochondria are shown. Scale bar = 10 μm. **c**, Population data and mean values comparing astrocyte mitochondrial Ca^2+^ event frequency (left), amplitude (middle) and half-width (right) (35 astrocytes and 5 MCU^−/−^ mice; n= 39 somatic, 99 branch, and 157 branchlet mitochondria, and 29 astrocytes from 4 WT littermates; n= 32 somatic, 148 branch, and 236 branchlet mitochondria). Errors are ± s.e.m. p-values are based on Mann-Whitney test. **d**, Representative time compressed confocal image of a DLS astrocyte during Ca^2+^ imaging with Ru265 application. Somatic, branch, and branchlet representative mitochondria are shown by magenta, orange, and blue circles, respectively. Scale bar = 15 μm. **e**, Representative Ca^2+^ event traces for somatic (magenta), branch (orange), and branchlet (blue) mitochondria with spontaneous Ca^2+^ events and after application of 1 μM Ru265 (left) and 10 μM Ru265 (right). **f**, Average Ca^2+^ event frequency of mitochondria before and after 1 μM Ru265 (left graph) (7 astrocytes and 4 mice; n= 11 somatic, 62 branch, and 80 branchlet mitochondria, and 10 μM Ru265, right graph from 7 astrocytes and 4 mice; n= 13 somatic, 57 branch, and 90 branchlet mitochondria). Errors are ± s.e.m. p-values for comparison of MCU^−/−^ and WT littermates are based on Mann-Whitney test. p-values for 1 μM or 10 μM Ru265 data are based on Wilcoxon Signed Rank test.

All drugs were bath perfused using a peristaltic pump (Harvard Apparatus) and the time taken for bath perfusion of drugs were set prior to each imaging session. Spontaneous activity in mitochondria was recorded for 5 min and drugs were sequentially applied with the order of application of drugs randomized for each slice. Concentrations of drugs applied were as follows: 300 μM glutamate, 5 μM SKF-38393, 10 μM quinpirole, 1 μM or 10 μM Ru265 (membrane permeable)^32^, and 20 μM cyclopiazonic acid (CPA). 1 μM Ru265 was introduced to the bath, followed by 10 μM Ru265 after a 10 min washout. For zero Ca^2+^ experiments CaCl_2_ was omitted from the recording buffer^16^. For CPA experiments, slices were incubated with 20 μM CPA for 15 min prior to imaging and CPA was maintained in the bath during imaging. Washout times for each drug were as follows: glutamate (10 min), SKF-38393 (35 min), quinpirole (35 min), zero Ca^2+^ (10 min), Ru265 (10 min), CPA (no washout possible).

### Data analyses

Ca^2+^ signals were analyzed using MiniAnalysis program 6.0.7 (Synaptosoft) and Origin (2019, Origin Lab Corp.). Image analyses were performed using ImageJ version 1.52 (NIH). Ca^2+^ imaging data were analyzed as previously described^16^. Briefly, image stacks were drift corrected in the x-y direction using the Turboreg plugin in ImageJ. 1-5% laser power at 633 nm caused significant photobleaching over the imaging period and this was corrected using EMBLtools (ImageJ plugin) with a frame-wise exponential fit.

Following drift and bleach correction MTDR labeled regions of interest (ROIs) were isolated using GECIquant as previously described^16^. We utilized an area constraint of 5-2000 μm^2^ for somatic and large territory mitochondria and 1-4 μm^2^ for small territory mitochondria. ROIs obtained in this way were used to extract Ca^2+^ events from GfaABC_1_D-mito-7-GCaMP6f labelled astrocytic mitochondria. ROIs containing no Ca^2+^ events were manually screened and eliminated. Ca^2+^ event frequency (events/min), amplitude (dF/F), and half-width (s) were analyzed using Minianalysis 6.0.07 (Synaptosoft).

### Statistical analyses

Ca^2+^ event measurements of frequency, amplitude and half widths were obtained from individually demarcated ROIs of mitochondria in the soma, branches or branchlets within individual astrocytes. Thus, each data point in the scatter plots represents one mitochondrial locus from one astrocyte. Averages for each of the Ca^2+^ event parameters were obtained from 3 – 4 astrocytes per slice, 2 – 4 slices per mouse, and between 4 – 17 mice per condition. Exact sample sizes for each experiment are specified in the figure legends. Slices from male and female mice were pooled together since we did not observe sex differences in our experiments. Statistics were performed using Origin Lab. To test for statistical significance, data were first tested for normality using Shapiro-Wilk test, and in the case of datasets with normally distributed data, either student’s unpaired t-test or paired t-test was used. Datasets containing non-normally distributed data were tested using Mann-Whitney (for unpaired data) or the Wilcoxon Signed Rank test (for paired data). Data were considered statistically significant if *p* < 0.05. Statistical tests used for each experiment used are indicated in figure legends.

## Results

### AAV 2/5 GfaABC_1_D-mito-7-GCaMP6f expresses GCaMP6f in astrocytic mitochondria

To specifically express GCaMP6f in astrocytic mitochondria, we generated an adeno-associated viral vector (AAV) with the astrocyte-specific GfaABC1D promoter^33^ driving expression of GCaMP6f tagged to a mitochondrial mito-7 targeting sequence (AAV 2/5 GfaABC_1_D-mito-7-GCaMP6f). AAV 2/5 GfaABC_1_D-mito-7-GCaMP6f virus was stereotaxically injected into the DLS of WT C57BL/6 mice. Three weeks later, live striatal slices were obtained from AAV-injected mice, labeled with MTDR, and imaged using a confocal microscope (Fig 1a).

MTDR co-localized with AAV-expressed GCaMP6f in discrete punctate structures within the soma, proximal primary branches, and peripheral branchlets of astrocytes (Fig. 1b). To further confirm co-localization of GCaMP6f in the mitochondria of astrocytes, we immunostained a separate set of DLS sections from AAV 2/5 GfaABC_1_D-mito-7-GCaMP6f-injected mice with the mitochondrial matrix protein pyruvate dehydrogenase (PDH)^34,35^. These sections showed co-localization of PDH with GFP antibody-labeled GCaMP6f in DLS astrocytes (Supplementary Fig. 1). Thus, using two independent methods, i.e. live imaging of striatal slices with MTDR, and immunostaining with mitochondria-specific PDH, we confirmed that AAV 2/5 GfaABC_1_D-mito-7-GCaMP6f specifically expressed GCaMP6f in astrocytic mitochondria.

### The size of functional mitochondria in DLS astrocytes depends on their subcellular location

We used the live DLS brain slices obtained from AAV 2/5 GfaABC_1_D-mito-7-GCaMP6f injected adult mice to demarcate punctate MTDR labeled ROIs that also co-express GCaMP6f in astrocytic mitochondria. Areas of GCaMP6f + MTDR labeled punctate structures were measured and segregated according to size. Spatially segregated mitochondria were observed in the somata, primary branches, and peripheral branchlets of all imaged astrocytes. Area analysis revealed significantly different sizes of mitochondria in the somata versus branches and branchlets of astrocytes (Fig. 1c). The largest mitochondria were somatic, with an average area of 21.6 ± 2.3 μm^2^. Mitochondria in astrocyte territories were significantly smaller than somatic mitochondria with average areas of 13.8 ± 0.9 μm^2^ for primary branch, and 1.7 ± 0.7 μm^2^ for peripheral branchlets (Fig. 1c). Thus, depending on their subcellular localization (somata versus territory), mitochondria in DLS astrocytes show clear variations in their size, with the largest mitochondria appearing in the somata, and the smallest ones in the most peripheral branchlets.

### Astrocytic mitochondria in the DLS show heterogenous spontaneous Ca^2+^ events

Live DLS slices from adult mice expressing mito-7-GCaMP6f were imaged for spontaneous Ca^2+^ events in mitochondria. Spontaneous Ca^2+^ events were observed in all three types of mitochondria (somatic, primary branches, and peripheral branchlets) (Fig. 1d and Supplementary movie 1). Average Ca^2+^ event frequencies in all three mitochondria populations were similar (1.1 ± 0.05 events/min), but interestingly, all mitochondria displayed a very discrete frequency distribution pattern, showing highly consistent increments of 0.25 events/min (Fig. 1e). In order to determine if discrete frequency patterns occurred because all mitochondria in a single astrocyte flux Ca^2+^ at a single specific frequency or if each astrocyte contained a mixture of mitochondrial Ca^2+^ event frequencies, we plotted the Ca^2+^ flux frequencies of individual mitochondria for each DLS astrocyte. We found that individual astrocytes display heterogenous mitochondrial Ca^2+^ event frequencies (Supplementary Fig. 2), suggesting a subcellular, rather than *en masse* regulation of astrocytic mitochondrial Ca^2+^ event frequencies in the DLS.

We found that amplitudes and half-widths of Ca^2+^ events in DLS astrocytes differed significantly among somatic, branch and branchlet mitochondria (Fig. 1e). Somatic mitochondria displayed the largest amplitude (0.93 ± 0.09 dF/F), followed by secondary branchlet (0.52 ± 0.01 dF/F) and primary branch mitochondria (0.41 ± 0.02 dF/F). Ca^2+^ events in somatic mitochondria also demonstrated the longest half-width (3.65 ± 0.17 s), followed by branch (2.79 ± 0.07 s) and branchlet mitochondria (1.01 ± 0.04 s). Thus, in addition to morphological heterogeneity, astrocytic mitochondria show specific differences in the kinetics of Ca^2+^ events, and this appears to be determined by the subcellular localization of mitochondria within an astrocyte.

### Ca^2+^ events in astrocytic mitochondria require endoplasmic reticulum (ER) Ca^2+^ stores

We next assessed potential sources for spontaneous Ca^2+^ events in DLS astrocytic mitochondria. To empty ER Ca^2+^ stores, live DLS slices were exposed for 15 min to 20 μM of the SERCA ATPase inhibitor, cyclopiazonic acid (CPA). We found that CPA caused a dramatic 4-fold decrease in Ca^2+^ event frequency for somatic, branch and branchlet mitochondria (Fig. 2a,b,d and Supplementary movie 2). Bath perfusion of slices with zero Ca^2+^ ACSF, however, did not alter mitochondrial Ca^2+^ events (Fig. 2c,e and Supplementary movie 3). The few remaining Ca^2+^ events after CPA showed significantly decreased amplitudes in all mitochondria (2.5-fold for soma and 1.5-fold for territory mitochondria), while half-widths remained largely unchanged (Supplementary Fig. 3). By contrast, zero Ca^2+^ ACSF had minimal effect on mitochondrial Ca^2+^ event amplitudes and half-widths (Supplementary Fig. 3). Based on these data, we conclude that the ER is a major source of Ca^2+^ fluxes in DLS astrocytic mitochondria, with very little contribution from extracellular calcium.

### Astrocytic mitochondria in the DLS do not flux Ca^2+^ through the mitochondrial calcium uniporter, MCU

Having found that mitochondrial Ca^2+^ events in DLS astrocytes primarily depend on ER stores, we sought to determine whether the mitochondrial calcium uniporter, MCU^36,37^ is a major portal for entry of Ca^2+^ into astrocytic mitochondria. We injected AAV 2/5 GfaABC_1_D-mito-7-GCaMP6f into the DLS of MCU^−/−^ mice in an outbred CD1 genetic background, which survives into adulthood despite the knockout of MCU^28^. Surprisingly, live DLS slices obtained from these mice displayed spontaneous astrocytic mitochondrial Ca^2+^ events that were indistinguishable from their WT littermates (Fig. 3a-c and Supplementary movie 4).

To address the possibility of compensatory mechanisms in CD1 MCU^−/−^ mice^38^, a selective and membrane permeable MCU blocker, Ru265^32^, was bath perfused onto AAV 2/5-GfaABC1D-mito-7-GCaMP6f injected DLS slices from WT C57Bl6 mice. Exposure to either 1 or 10 μM Ru265 did not inhibit mitochondrial Ca^2+^ event amplitudes or half-widths (Supplementary Fig. 4), but both concentrations of Ru265 caused a 25% decrease in the inter-frequency interval for all mitochondrial subpopulations from 0.25 to 0.1 event/min (Fig. 3d-f). These data suggest that rather than being the primary portal for Ca^2+^ flux in astrocytic mitochondria, MCU likely regulates the frequency of mitochondrial Ca^2+^ events in astrocytes.

### Ca^2+^ events in mitochondria of DLS astrocytes are distinct from those in the HPC

Astrocytes in the DLS possess a significantly different proteomic and transcriptional profile from the HPC^39^, and astrocyte populations have been shown to be as heterogenous as neurons^40,41^. Based on these findings, we asked if astrocytic mitochondria from these two brain regions also display heterogeneity. We compared the morphological profile and spontaneous Ca^2+^ event kinetics of astrocytic mitochondria from the DLS with those in the HPC.

The CA1 region in the HPC of WT C57BL/6 mice was stereotaxically injected with AAV2/5 GfaABC_1_D-mito-7-GCaMP6f. Live HPC slices from these mice were labeled with MTDR and imaged with a confocal microscope. Similar to DLS astrocytes, HPC astrocytes showed robust GCaMP6f expression that co-localized with MTDR to the soma, primary branches, and peripheral branchlets (Fig. 4a). Interestingly, somatic mitochondria in HPC astrocytes were 2-fold smaller than DLS astrocytes, while mitochondria in branches and branchlets were of similar size to DLS astrocytes (Fig. 4b).

**Figure 4.**
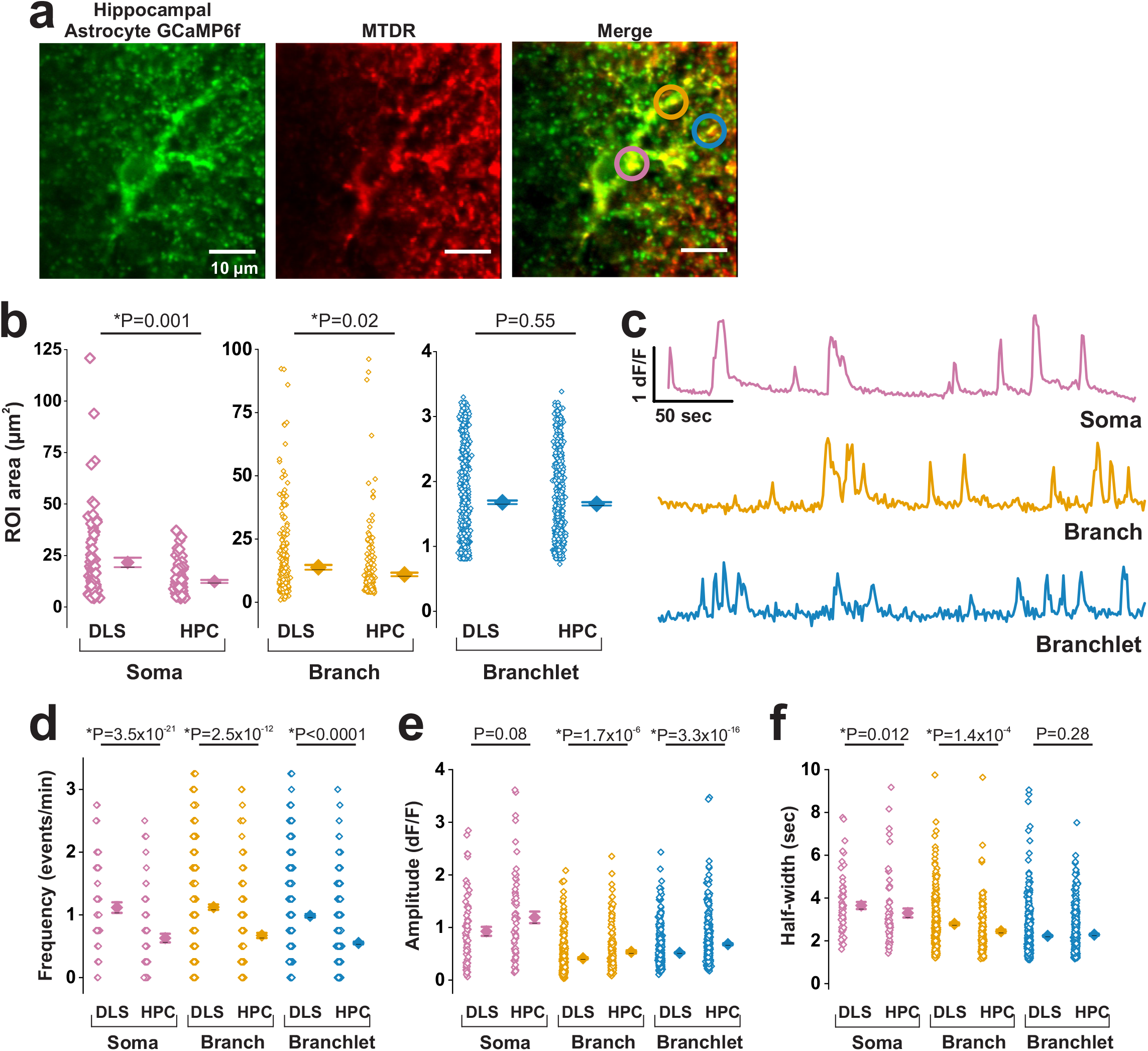
DLS and HPC astrocytes differ in mitochondrial sizes and Ca^2+^ event frequencies. **a**, Representative time compressed confocal images of a hippocampal (HPC) astrocyte from the CA1 region expressing GCaMP6f (green) + MTDR (red). Somatic, branch, and branchlet mitochondria are indicated by magenta, orange, and blue circles respectively. Scale bar = 10 μm. **b**, Population data and mean values comparing mitochondrial areas between 56 DLS astrocytes from 15 mice (n= 67 somatic, 336 branch, and 605 branchlet mitochondria) and 55 HPC astrocytes from 8 mice (n= 79 somatic, 211 branch, and 547 branchlet mitochondria). **c**, Representative traces of spontaneous mitochondrial Ca^2+^ events in somatic (magenta), branch (orange), and branchlet (blue) mitochondria of HPC astrocytes. **d-f**, Comparison of spontaneous astrocytic mitochondrial Ca^2+^ event frequency, amplitude and half-width between DLS and HPC astrocytes from 56 DLS astrocytes and 15 mice (n= 67 somatic, 336 branch, and 605 branchlet mitochondria), and from 55 HPC astrocytes and 8 mice (n= 79 somatic, 211 branch, and 547 branchlet mitochondria). Errors are ± s.e.m. p-values are based on Mann-Whitney test.

We found that astrocytic mitochondria in the HPC showed spontaneous Ca^2+^ events (Fig. 4c and Supplementary movie 6). Irrespective of their localization to somata, branches or branchlets, all astrocytic mitochondria in the HPC displayed Ca^2+^ event frequencies that were half the frequency of Ca^2+^ events in DLS astrocytes (0.6 events/min for HPC versus 1.1 events/min for DLS) (Fig. 4d). Despite a lower average frequency of Ca^2+^ events in HPC astrocytic mitochondria, the interval between Ca^2+^ events in the HPC was always 0.25 event/min, which was similar to the DLS. By contrast, we found that amplitudes of Ca^2+^ events in HPC mitochondria were generally larger than those in the DLS for somatic, branch and branchlet mitochondria (Fig. 4e), but Ca^2+^ event half-widths were similar for both regions (Fig. 4f). Together, these data show that astrocytic mitochondria in the DLS differ from the HPC with regard to morphology, as well as the frequencies and amplitudes of Ca^2+^ fluxes.

### Mitochondrial Ca^2+^ events in DLS and HPC astrocytes show dual responses to glutamate

Since the DLS and HPC receive glutamatergic input from the cortex^42–45^, we assessed the effects of glutamate on astrocytic mitochondrial Ca^2+^ events in both these brain regions.

Amplitudes and half-widths of astrocyte mitochondrial Ca^2+^ events in the DLS remained unchanged with bath application of 300 μM glutamate (Supplementary Fig. 5a,b). However, glutamate exposure resulted in a dual effect on astrocytic mitochondrial Ca^2+^ event frequency. Bath application of 300 μM glutamate decreased the Ca^2+^ event frequency by 43 ± 10% in somatic, 46 ± 7% in branch, and 54 ± 5% in branchlet mitochondria, while other mitochondria within the same astrocytes showed an increase in event frequency by 57 ± 10% in somatic, 54 ± 7% in branch, and 46 ± 5% in branchlet mitochondria (Figs. 5a-c). Thus, each DLS astrocyte displayed a mixture of decreased or increased mitochondrial Ca^2+^ event frequency. For both effects of glutamate, *viz.* a decrease or increase in mitochondrial Ca^2+^ event frequency, glutamate invariably increased the dynamic range of mitochondrial Ca^2+^ event frequencies (Fig. 5c).

**Figure 5.**
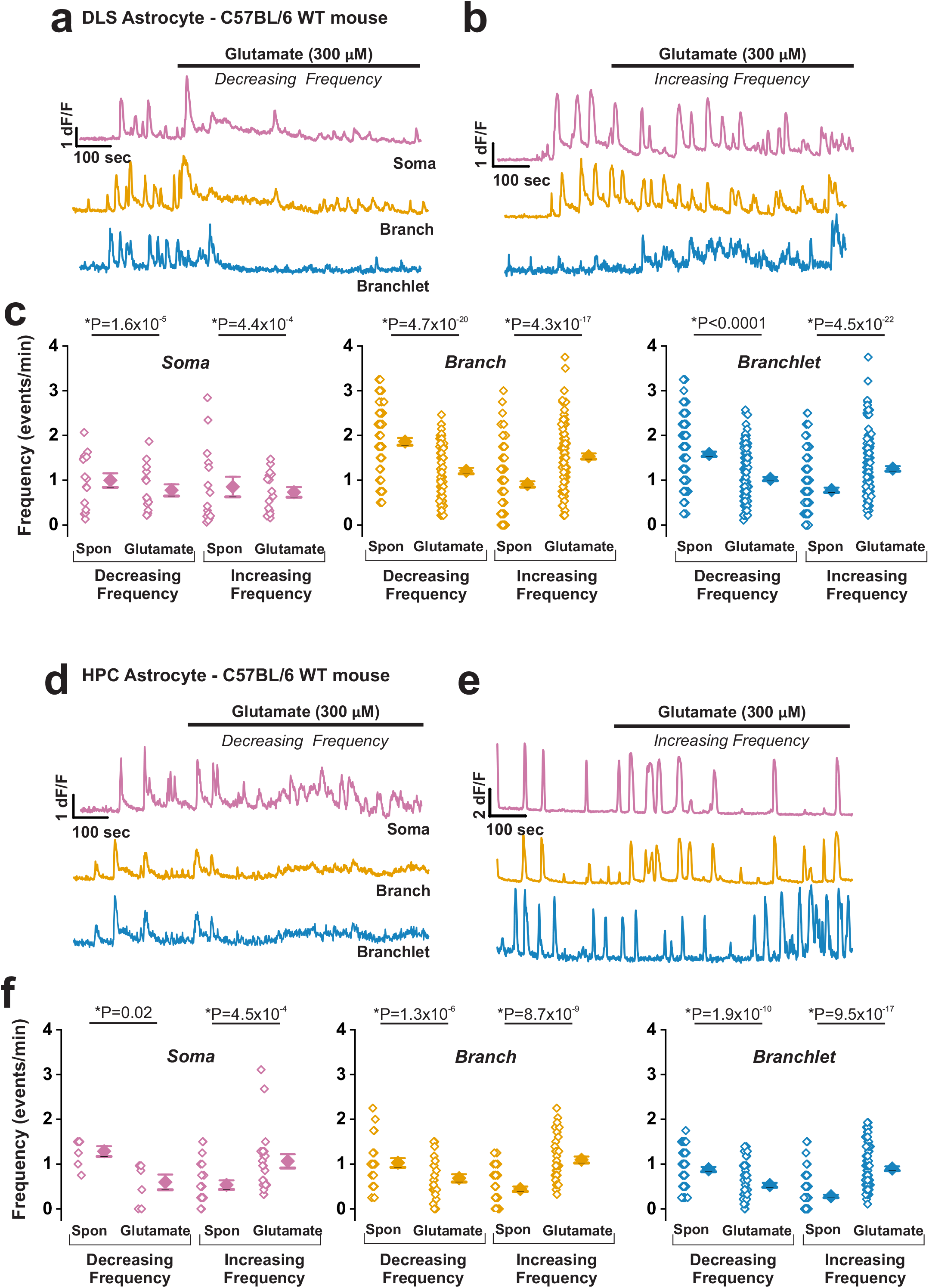
Glutamate alters mitochondrial Ca^2+^ event frequencies in DLS and HPC astrocytes. Representative Ca^2+^ traces for somatic (magenta), branch (orange), and branchlet (blue) mitochondria from DLS astrocytes with either **a**, decreasing or **b**, increasing Ca^2+^ event frequency in response to glutamate. **c**, Population data and mean values for DLS mitochondrial Ca^2+^ event frequency from 23 DLS astrocytes and 12 mice. Mitochondria in graphs are segregated based on whether glutamate decreased or increased Ca^2+^ event frequency (n= 15 decreasing and 16 increasing somatic, 75 decreasing and 110 increasing branch, and 165 decreasing and 135 increasing branchlet mitochondria). **d-e**, As in **a-b**, but for HPC astrocytic mitochondria in response to glutamate. **f**, As in **c**, but for HPC mitochondria from 17 astrocytes and 8 mice (n= 7 decreasing and 21 increasing somatic, 25 decreasing and 44 increasing branch, and 59 decreasing and 97 increasing branchlet mitochondria). Errors are ± s.e.m. p-values are based on paired sample t-test for decreasing and increasing frequency in DLS somatic mitochondria and decreasing frequency of DLS branch mitochondria. All other DLS data sets are based on Wilcoxon Signed Rank test. For data from HPC astrocytes, p-values are based on a paired t-test for decreasing frequency in HPC branch mitochondria, while all other data sets are based on Wilcoxon Signed Rank test.

Similar to the DLS, bath application of glutamate to HPC slices did not alter mitochondrial Ca^2+^ event amplitudes and half-widths (Supplementary Fig. 5c,d), but caused a dual response in mitochondrial Ca^2+^ event frequency for all astrocytes that were imaged. For the HPC, we observed a decrease in frequency by 21 ± 10% in somatic, 46 ± 11% in branch, and 50 ± 9% in branchlet mitochondria and an increase in frequency by 79 ± 10% in somatic, 52 ± 11% in branch, and 50 ± 9% in branchlet mitochondria (Fig. 5d-f). Similar to the DLS, glutamate eliminated the regular frequency spacing in all HPC astrocytic mitochondria.

### Mitochondrial Ca^2+^ events in DLS and HPC astrocytes show dual responses to dopaminergic D1 and D2 receptor agonists

Dopaminergic neurons in the substantia nigra pars compacta (SNc) project to DLS and the HPC ^46,47^. Astrocytes in the DLS and HPC are therefore constantly exposed to dopamine *in vivo*, which would result in a sustained activation of D1 and D2 dopamine receptors in both brain regions. We assessed the effects of the D1–specific agonist, SKF-38393 and the D2 –specific agonist, quinpirole on mitochondrial Ca^2+^ events in astrocytes from the DLS and HPC. Similar to glutamate, bath application of 5 μM SKF-38393 and 10 μM quinpirole induced dual effects on Ca^2+^ event frequencies in somatic, branch, and branchlet mitochondria of DLS and HPC astrocytes (Figs. 6 and 7).

**Figure 6.**
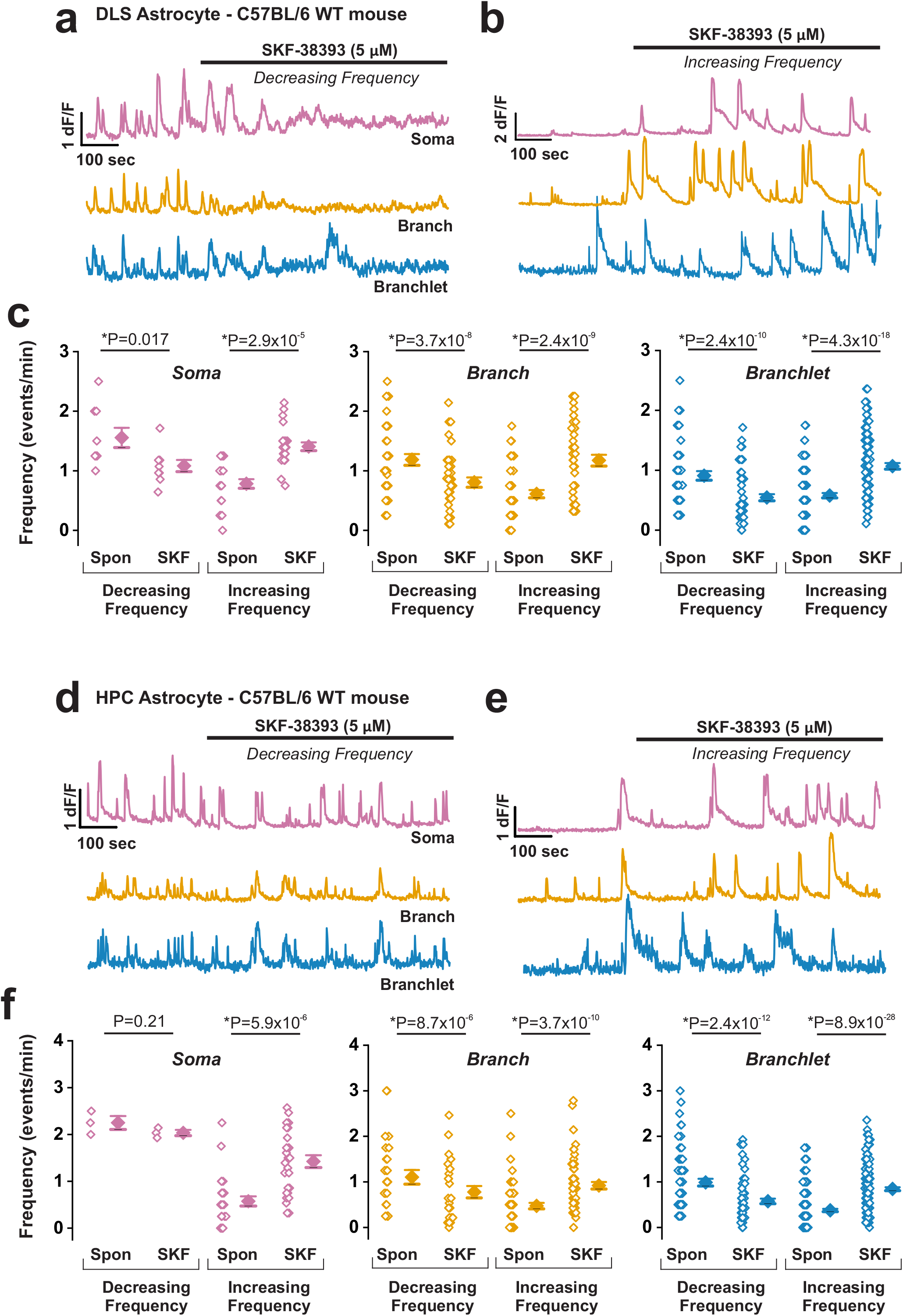
Responses of mitochondrial Ca^2+^ event frequency in DLS and HPC astrocytes to the D1-receptor agonist, SKF-38393. Representative Ca^2+^ traces for somatic (magenta), branch (orange), and branchlet (blue) mitochondria **a**, decreasing or **b**, increasing Ca^2+^ event frequency in response to SKF-38393. **c**, Population data and mean values for mitochondrial Ca^2+^ event frequency from 17 DLS astrocytes and 8 mice (n= 9 decreasing and 23 increasing somatic, 40 decreasing and 47 increasing branch, and 53 decreasing and 105 increasing branchlet mitochondria) before and after SKF-38393. **d-e**, As in **a-b**, but for HPC astrocytic mitochondria with SKF-38393 application. **f**, As in **c**, but from mitochondria in 17 HPC astrocytes and 7 mice (n= 3 decreasing and 27 increasing somatic, 26 decreasing and 57 increasing branch, and 65 decreasing and 165 increasing branchlet mitochondria) before and after SKF-38393. Errors are ± s.e.m. p-values are based on paired sample t-test for decreasing frequency in DLS somatic mitochondria. All other data sets for DLS and HPC astrocytes are based on Wilcoxon Signed Rank test.

**Figure 7.**
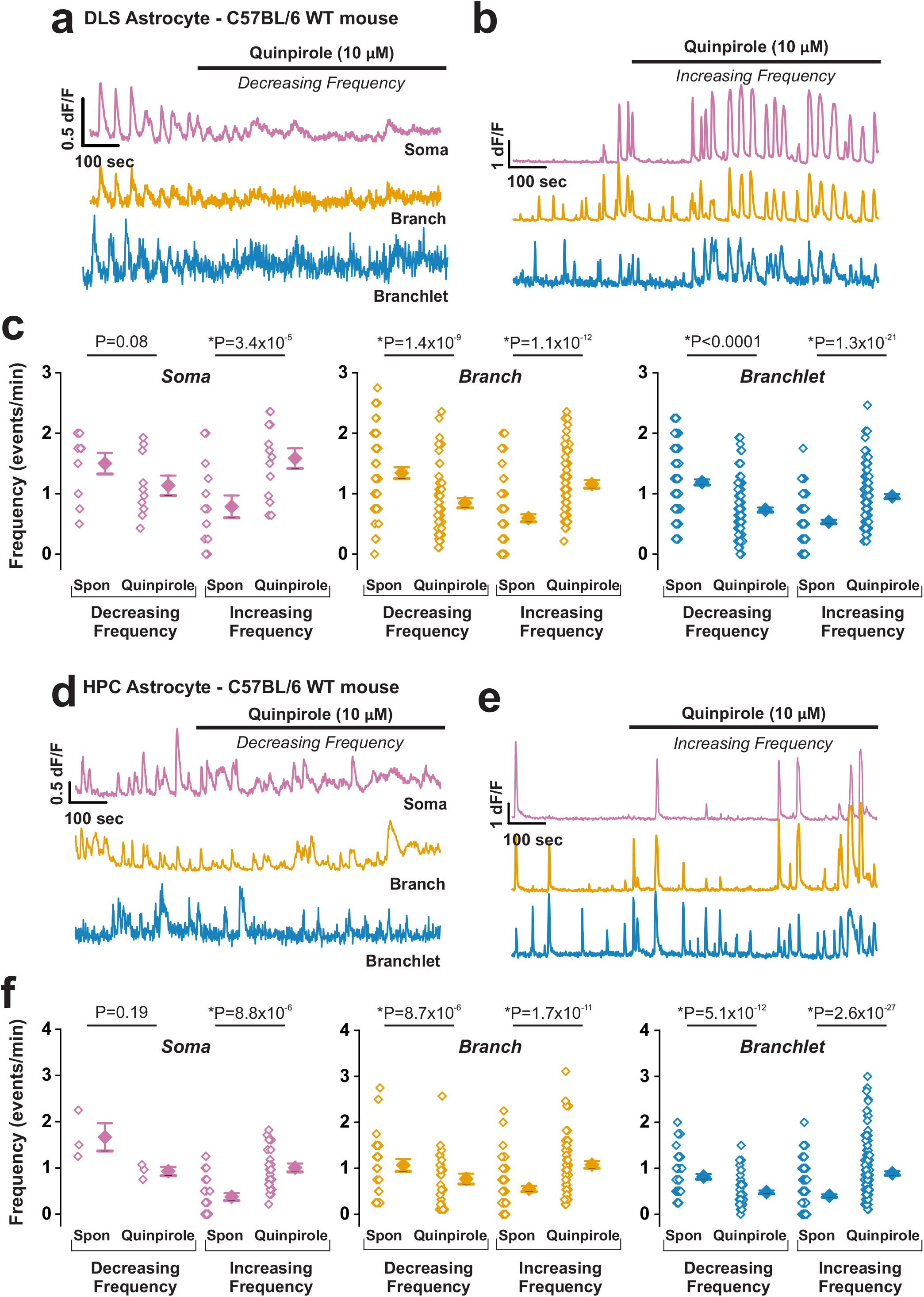
Responses of mitochondrial Ca^2+^ event frequency in DLS and HPC astrocytes to the D2-receptor agonist, quinpirole. Representative Ca^2+^ event traces for somatic (magenta), branch (orange), and branchlet (blue) mitochondria in DLS astrocytes in response to quinpirole **a**, decreasing or **b**, increasing Ca^2+^ event frequency. **c**, Population data and mean values for DLS mitochondrial Ca^2+^ event frequency changes in response to quinpirole from 21 DLS astrocytes and 11 mice (n= 10 decreasing and 14 increasing somatic, 51 decreasing and 67 increasing branch, and 129 decreasing and 121 increasing branchlet mitochondria). **d-e**, As in **a-b**, but for mitochondria from HPC astrocytes. **f**, As in **c**, but for mitochondria from 16 HPC astrocytes and 7 mice (n= 3 decreasing and 26 increasing somatic, 26 decreasing and 60 increasing branch, and 63 decreasing and 157 increasing branchlet mitochondria). Errors are ± s.e.m. p-values are based on paired sample t-test for decreasing and increasing frequency in DLS somatic mitochondria and decreasing frequency in HPC somatic mitochondria. All other data sets for DLS and HPC astrocytes are based on Wilcoxon Signed Rank test.

For the DLS, the D1-specific agonist, SKF-38393 decreased Ca^2+^ event frequencies by 29 ± 9% in somatic, 49 ± 8% in primary branches, and 38 ± 6% in secondary branchlet mitochondria and increased Ca^2+^ event frequencies by 71 ± 9% in somatic, 51 ± 8% in primary branches, and 61 ± 6% in secondary branchlet mitochondria of astrocytes (Fig. 6a-c). Astrocytic mitochondria in the HPC also displayed a dual response to SKF-38393 (Fig. 6d-f). SKF-38393 exposure decreased mitochondrial Ca^2+^ events in HPC astrocytes by 15 ± 8% in somatic, 32 ± 7% in primary branches, and 30 ± 4% in secondary branchlets, and increased mitochondrial Ca^2+^ events by 83 ± 8% in somatic, 68 ± 7% in primary branches, and 70 ± 4% in secondary branchlets.

The D2-specific agonist, quinpirole decreased Ca^2+^ event frequencies in DLS astrocytic mitochondria by 45 ± 11% in somatic, 51 ± 8% in primary branches, and 50 ± 5% in secondary branchlet mitochondria, but increased Ca^2+^ event frequencies in other mitochondria by 55 ± 11% in somatic, 49 ± 8% in primary branches, and 52 ± 5% in secondary branchlets (Figs. 7a-c). In the HPC, quinpirole caused a decrease in mitochondrial Ca^2+^ events by 11 ± 8% in somatic, 37 ± 7% in primary branches, and 30 ± 5% in secondary branchlets and an increase in Ca^2+^ events of other mitochondria from the same HPC astrocytes by 89 ± 8% in somatic, 63 ± 7% in primary branches, and 70 ± 5% in secondary branchlets (Fig. 7d-f). Neither SKF-38393 nor quinpirole altered amplitudes and half-widths of mitochondrial Ca^2+^ events in either the DLS or the HPC (Supplementary Figs. 6,7). However, regardless of whether they caused a decrease or an increase in mitochondrial Ca^2+^ event frequencies, both SKF-38393 and quinpirole dramatically increased the dynamic range of Ca^2+^ event frequencies in all astrocytic mitochondria from the DLS and HPC (Figs. 6c and 7c).

In summary, these data show that activating either glutamate or D1 and D2 neurotransmitter receptors in DLS and HPC astrocytes causes a dual response in mitochondrial Ca^2+^ event frequency. Thus, while some mitochondria show a decrease in frequency, other mitochondria in the same astrocyte robustly increase Ca^2+^ event frequencies.

### Mitochondrial Ca^2+^ responses to neurotransmitter receptor agonists do not depend on MCU

We assessed the effect of glutamate, SKF-38393, and quinpirole on the mitochondrial Ca^2+^ event frequency in DLS astrocytes from MCU^−/−^ mice. Following exposure to all three agonists (glutamate, SKF-38393, and quinpirole), astrocytes from MCU^−/−^ mice displayed dual responses of mitochondrial Ca^2+^ event frequencies and dramatic changes in frequency distributions of mitochondrial Ca^2+^ events that were indistinguishable from WT littermate controls (Figs. 8-10). These data suggest that MCU does not play a role in neurotransmitter-induced responses of astrocytic mitochondrial Ca^2+^ events.

**Figure 8.**
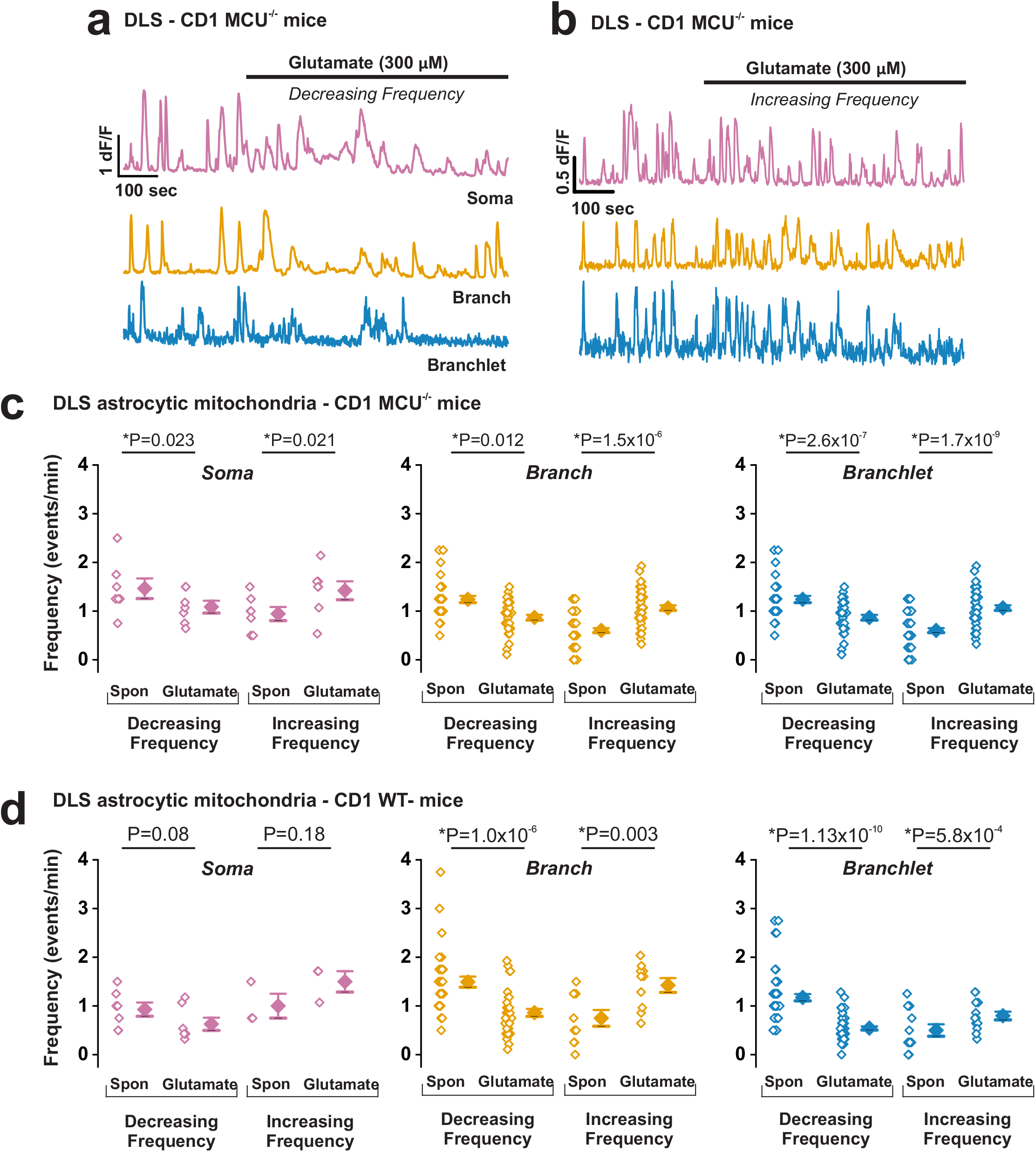
DLS astrocytic mitochondrial Ca^2+^ event responses to glutamate are not altered in MCU^−/−^ mice. Representative Ca^2+^ event traces for soma (magenta), branch (orange), and branchlet (blue) mitochondria from DLS astrocytes of MCU^−/−^ mice showing either **a**, decreasing or **b,** increasing Ca^2+^ event frequency in response to glutamate application. **c**, Population data and mean values for spontaneous and glutamate induced Ca^2+^ event frequency from 10 DLS astrocytes in 5 MCU^−/−^ mice (n= 7 decreasing and 10 increasing somatic, 18 decreasing and 18 increasing branch, and 35 decreasing and 48 increasing branchlet mitochondria). **d**, As in **c**, but for mitochondria from 10 DLS astrocytes in 4 WT littermates (n= 3 decreasing and 10 increasing somatic, 34 decreasing and 10 increasing branch, and 56 decreasing and 13 increasing branchlet mitochondria). Errors are ± s.e.m. p-values for MCU^−/−^ mice are based on paired sample t-test except that branchlet mitochondria were subject to Wilcoxon Signed Rank test. p-values for WT littermates are based on paired t-tests, except decreasing frequency branch mitochondria and decreasing frequency branchlet mitochondria which were subject to Wilcoxon Signed Rank test.

## Discussion

We generated an AAV construct, AAV2/5 GfaABC1D-mito-7-GCaMP6f, to measure the influx of Ca^2+^ into astrocytic mitochondria from DLS and HPC brain slices. Our experiments reveal several unique features in astrocytic mitochondria, such as: (i) Significant differences in mitochondrial size based on subcellular location, as well as the brain region in which the astrocyte resides, (ii) Divergence of spontaneous Ca^2+^ event kinetics in somatic, branch, and branchlet mitochondria, (iii) Strong dependence of mitochondrial Ca^2+^ fluxes on ER Ca^2+^ stores, (iv) The surprising lack of a major role for the mitochondrial calcium uniporter, MCU in Ca^2+^ fluxes, and (v) Dual responses of Ca^2+^ event frequencies to multiple neurotransmitter agonists. Taken together, these findings provide a comprehensive foundational understanding of mitochondrial Ca^2+^ fluxes in astrocytes within disease-relevant regions of the brain.

Using AAV-expressed GfaABC1D-mito-7-GCaMP6f co-labeled with MTDR, we found that DLS and HPC astrocytes possess distinct sizes of mitochondria in their soma, branches, and branchlets. The largest mitochondria localized to astrocytic soma, while the smallest ones were observed in peripheral secondary branchlets, and intermediate size mitochondria localized to primary branches of astrocyte territories (Figs. 1c and 4b). This finding is in agreement with previous studies showing similar distributions of mitochondria in cortical astrocytes^23,48^. Notably, somatic mitochondria in HPC astrocytes were half the size of those in the soma of DLS astrocytes (Fig. 4b), which likely represents a functional rather than a morphological adaptation. Indeed, despite having similar somatic sizes, the number of neuronal cell bodies contacted by astrocytes in the DLS is greater than the CA1 region in the HPC^39^. Thus, somatic mitochondria in astrocytes within these two brain regions might cater to the specific energy requirements of neuronal somata rather than individual synapses, while mitochondria in branches and branchlets likely respond to activity within individual synapses.

All of the GCaMP6f-expressing mitochondria in DLS and HPC astrocytes displayed robust spontaneous Ca^2+^ events. When compared to Ca^2+^ events in somata, we found that branch and branchlet mitochondria showed significantly smaller amplitudes and shorter half-widths (Figs. 1e and 4d-f). In addition, we observed a highly structured frequency distribution of spontaneous mitochondrial Ca^2+^ events in DLS and HPC astrocytes (Fig. 4d). For each of the mitochondrial loci analyzed, we found that Ca^2+^ events occurred at a single frequency not determined by subcellular location, and when analyzed in the context of individual astrocytes, mitochondrial Ca^2+^ events displayed a mixture of frequencies rather than one uniform frequency. Because mitochondrial Ca^2+^ is a primary determinant of oxidative phosphorylation and the ability to generate ATP^49–51^, we suggest that there is a quantitative correlation between subcellular ATP generation in astrocytes and the magnitude of Ca^2+^ events in the soma versus territories. Thus, unlike the heterogenous mitochondrial Ca^2+^ events in astrocyte territories, which would cater to small scale synaptic activity, the larger, more invariant Ca^2+^ events in somatic mitochondria generate more ATP for regulating the baseline activity of multiple neurons in close proximity to any given astrocyte. Apart from these subcellular differences, we also observed region-specific differences in the kinetics of spontaneous astrocytic mitochondrial Ca^2+^ events. Mitochondria in all sub-compartments of HPC astrocytes fluxed Ca^2+^ at a 2-fold lower frequency than astrocytes in the DLS (Fig. 4d). Our simplest interpretation for this finding is that when compared to the DLS, neurons in the hippocampus possess lower levels of baseline activity, which would correlate with the amount of astrocyte-generated ATP, and the magnitude of mitochondrial Ca^2+^ events in astrocytes from each of these brain structures.

We did not directly assess the directionality of spontaneous astrocytic mitochondrial Ca^2+^ fluxes. However, because the mito-7 signaling sequence would cause GCaMP6f to localize to the inner mitochondrial matrix, one can assume that GCaMP6f detects an influx of Ca^2+^ into the inner mitochondrial matrix, rather than the efflux of Ca^2+^ from mitochondria. To determine subcellular sources for Ca^2+^ influx into mitochondria in DLS astrocytes, we sequentially depleted extracellular and ER Ca^2+^ stores. Depleting extracellular Ca^2+^ did not alter mitochondrial Ca^2+^ events (Fig. 2c,e). By contrast, emptying ER Ca^2+^ stores with the SERCA ATPase pump inhibitor, CPA almost completely abolished Ca^2+^ events (Fig. 2b,d), suggesting that Ca^2+^ stores in the ER serve as a major source for spontaneous Ca^2+^influx into astrocytic mitochondria. Calcium transfer from the ER to mitochondria likely occurs via physical domains, known as mitochondria-associated membranes (MAMs), which physically and functionally link the ER with mitochondria^52,53^. Strong evidence for the presence of MAMs in astrocytes comes from a recent study showing that astrocytic mitochondria extensively contact the ER through mitofusin tethering domains^54^. Although MAMs are enriched in ER-localized inositol triphosphate type 2 (IP3R2) receptors^55^, a previous study has shown that astrocytic mitochondrial Ca^2+^ fluxes are independent of IP3R2^23^. An alternative possibility is that Ca^2+^ transfer from the ER to mitochondria occurs through other isoforms of IP3 receptors, and ER-localized ryanodine receptors (RyRs)^56,57^. Thus, we surmise that spontaneous Ca^2+^ events in astrocytic mitochondria occur at MAMs, and most likely involve Ca^2+^ transfer to mitochondria via ER-localized isoforms of IP3 and RyR receptors.

Having confirmed the ER as a primary source of Ca^2+^ for astrocytic mitochondria, we sought to determine the extent to which the mitochondrial calcium uniporter, MCU plays a role for Ca^2+^ entry into the inner mitochondrial matrix. To this end, we examined mitochondrial Ca^2+^ events in astrocytes from MCU^−/−^ mice, and much to our surprise, found that Ca^2+^ events in DLS astrocytes from MCU^−/−^ mice were indistinguishable from WT littermates (Fig. 3a-c). Compensatory mechanisms in MCU^−/−^ mice are unlikely to play a role because exposure to the MCU-specific inhibitor, Ru265, failed to inhibit mitochondrial Ca^2+^ fluxes in DLS astrocytes from WT mice (Fig. 3d-f). We observed that the Ru265-mediated inhibition of MCU significantly altered the frequency distribution of Ca^2+^ events, which suggests that rather than acting as a major portal for Ca^2+^ entry into mitochondria, as observed in other cells^36^, MCU in astrocytes may regulate specific aspects of mitochondrial Ca^2+^ flux kinetics. Our data further indicate that the regulatory role of MCU may be restricted to spontaneous Ca^2+^ events in mitochondria, because when compared to WT littermates, exposure to neurotransmitter agonists did not alter mitochondrial Ca^2+^ fluxes in astrocytes from MCU^−/−^ mice. In the absence of MCU as a major portal for Ca^2+^ influx into the matrix of astrocytic mitochondria, one candidate molecule to consider is the mitochondrial sodium calcium exchanger, NCLX. Evidence for an important role of NCLX comes from a study showing that NCLX mediates spontaneous Ca^2+^ oscillations in mitochondria by sequentially alternating between forward, Ca^2+^ efflux and reverse Ca^2+^ influx transport modes^58^. In addition, siRNA-mediated or pharmacological inhibition of NCLX alters cytosolic Ca^2+^ fluxes and ATP responses in astrocytes^59^. These previous studies, along with our results strongly indicate that rather than MCU, the mitochondrial transporter, NCLX may be the primary portal for Ca^2+^ influx into the inner matrix of astrocytic mitochondria.

Finally, we assessed the response of mitochondrial Ca^2+^ events in astrocytes to neurotransmitters by bath perfusing DLS and HPC brain slices with glutamate, the D1 receptor-specific agonist SKF-38393, or the D2 receptor-specific agonist quinpirole. Each of these neurotransmitter receptor agonists caused dual responses in mitochondrial Ca^2+^ flux frequencies within the somata, branches, and branchlets of DLS and HPC astrocytes (Figs. 5-10). Specifically, while a proportion of astrocytic mitochondria increased their frequency, Ca^2+^ event frequencies in other mitochondria from the same astrocyte were significantly reduced. Although mitochondrial Ca^2+^ event frequencies were altered by all tested neurotransmitter receptor agonists, their amplitudes and half widths remained largely unchanged (Supplementary Figs. 5-7), which suggests that Ca^2+^ flux frequencies in mitochondria are a primary determinant of astrocyte responses to neurotransmitters. In line with our finding, a previous study utilized bulk loaded Fluo-3 AM into HPC astrocytes *in situ*, and showed that dopamine causes a similar dual response in astrocytic Ca^2+^ signals^60^. Here, we show that these previously reported dopamine-induced dual responses are in fact mediated by changes in the frequency of Ca^2+^ events in the mitochondria of astrocytes. Conceivably, Ca^2+^ event frequencies for any one mitochondrion in an astrocyte territory are likely to be synchronized with one or more associated microcircuits. This can cause neurotransmitters to independently alter Ca^2+^ event frequencies, based on energy requirements for a particular microcircuit linked to one or more astrocytic mitochondria. Divergent neurotransmitter-induced changes in the frequency of mitochondrial Ca^2+^ events within the same astrocyte would not only modulate ATP generation in microcircuits, but can also cause genetic and epigenetic changes in the astrocytes themselves^61^. Thus, responses of astrocytic mitochondrial Ca^2+^ fluxes to sustained neurotransmitter release will, in the long-term, alter signaling mechanisms between astrocytes and neurons.

**Figure 9.**
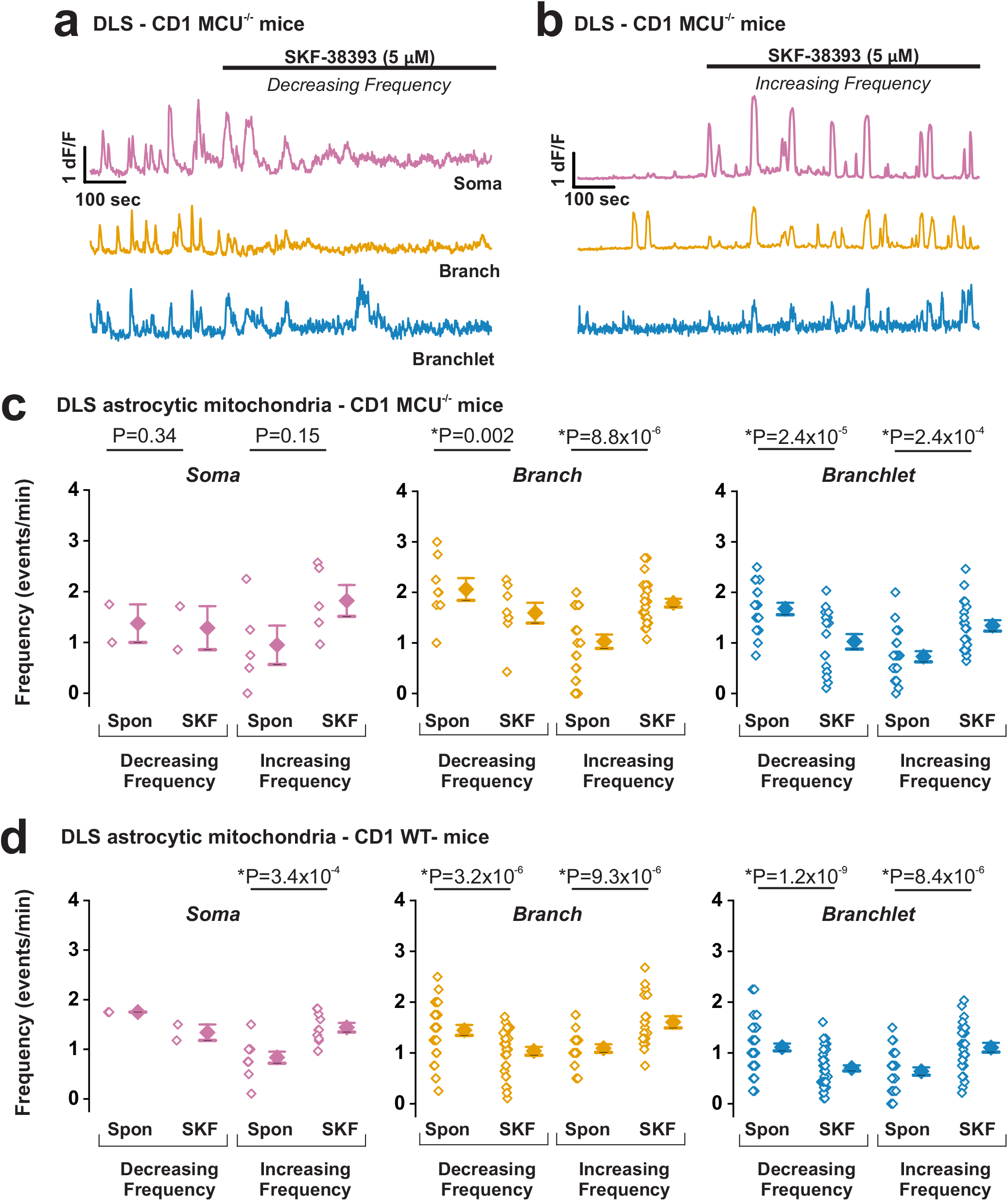
DLS astrocytic mitochondrial Ca^2+^ event responses to SKF-38393 are not altered in MCU^−/−^ mice. Representative Ca^2+^ traces for somatic (magenta), branch (orange), and branchlet (blue) mitochondria from DLS astrocytes with either **a**, decreasing or **b,** increasing Ca^2+^ event frequency in response to SKF-38393 application. **c**, Population data and mean values for spontaneous and SKF-38393 induced mitochondrial Ca^2+^ event frequency from 9 DLS astrocytes in 5 MCU^−/−^ mice (n= 2 decreasing and 5 increasing somatic, 8 decreasing and 26 increasing branch, and 17 decreasing and 21 increasing branchlet mitochondria). **d**, As in **c**, but for mitochondria from 9 DLS astrocytes in 4 WT littermates (n= 2 decreasing and 10 increasing somatic, 29 decreasing and 19 increasing branch, and 42 decreasing and 27 increasing branchlet mitochondria). Errors are ± s.e.m. p-values for MCU^−/−^ mice are based on paired t-tests except for increasing frequency branch mitochondria which is based on Wilcoxon Signed Rank test. All p-values for WT littermates are based on paired t-test.

**Figure 10.**
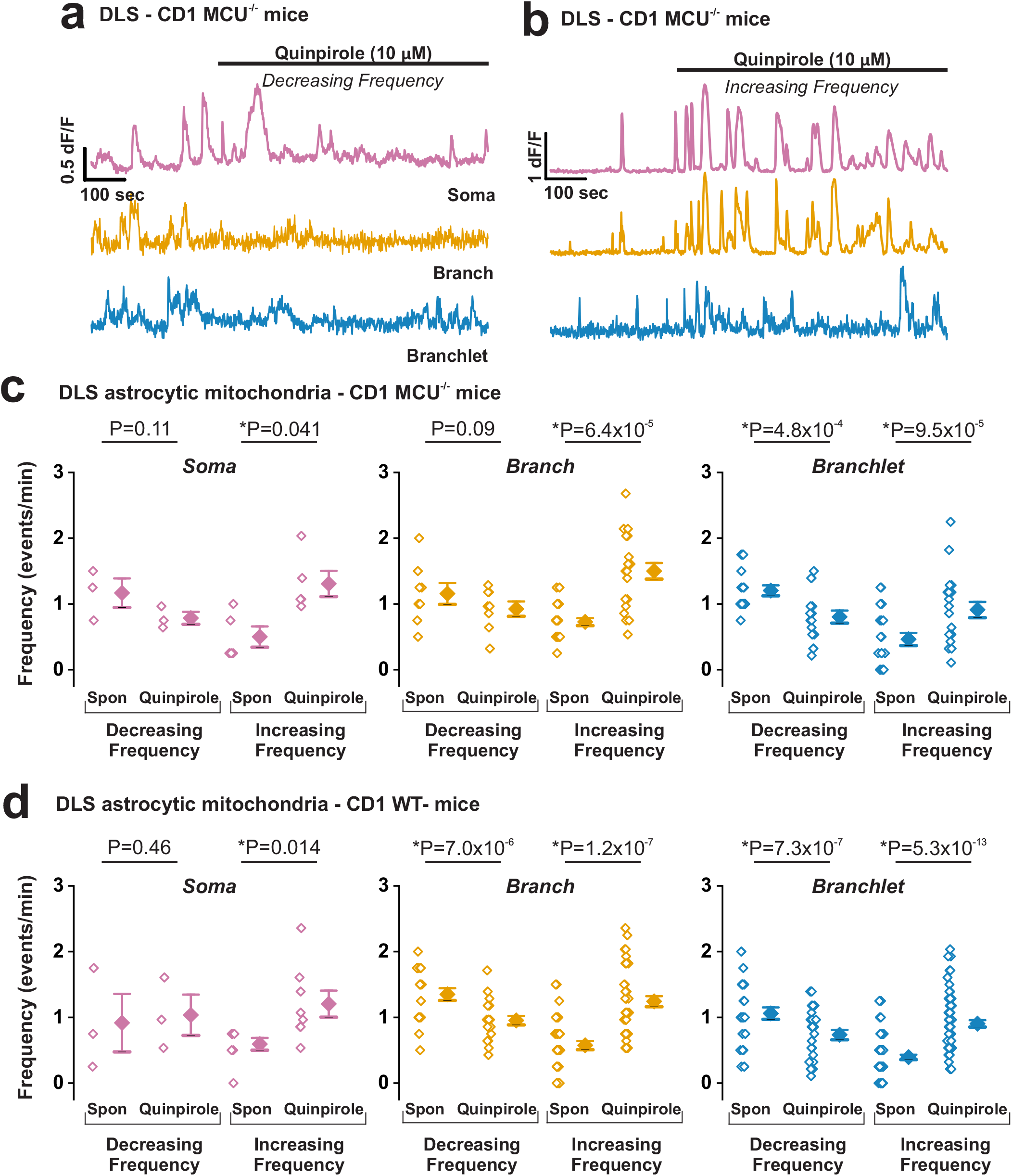
DLS astrocytic mitochondrial Ca^2+^ event responses to quinpirole are not altered in MCU^−/−^ mice. Representative Ca^2+^ traces for somatic (magenta), branch (orange), and branchlet (blue) mitochondria from DLS astrocytes with either **a**, decreasing or **b,** increasing Ca^2+^ event frequency in response to quinpirole. **c**, Population data and mean values for changes in quinpirole induced Ca^2+^ event frequency from 9 DLS astrocytes in 5 MCU^−/−^ mice (n= 3 decreasing and 5 increasing somatic, 8 decreasing and 21 increasing branch, and 16 decreasing and 20 increasing branchlet mitochondria). **d**, As in **c**, but for mitochondria from 10 DLS astrocytes in 4 WT littermates (n= 3 decreasing and 10 increasing somatic, 20 decreasing and 37 increasing branch, and 29 decreasing and 69 increasing branchlet mitochondria). Errors are ± s.e.m. p-values for MCU^−/−^ mice are based on paired sample t-test, except for branch mitochondria with increased in frequency and branchlet mitochondria with decreased or increased in frequency, that were subject to Wilcoxon Signed Rank test. p-values for WT littermates are based on paired sample t-test, except for increasing frequency branch mitochondria and increasing frequency branchlet mitochondria which were subject to Wilcoxon Signed Rank test.

In conclusion, our study shows that astrocytic mitochondria possess a unique physiological profile in the form of spontaneous Ca^2+^ fluxes, with significant differences from other cells in their kinetics and portals of Ca^2+^ entry. We also report subcellular and inter-regional differences, as well as dual responses of astrocytic mitochondrial Ca^2+^ fluxes to neurotransmitter agonists. Together, these results provide a strong mechanistic foundation for understanding how Ca^2+^ fluxes in the mitochondria of astrocytes change during neurodegeneration, stroke, or aging. We predict that our findings will enable specific manipulations of Ca^2+^ fluxes in astrocytic mitochondria as an effective treatment strategy for multiple neurological conditions.

## Acknowledgements

Supported by a pre-doctoral award from the National Science Foundation Graduate Research Fellowships Program (NSF-GRFP) to TEH and a grant from the Texas A&M University (TAMU) Clinical Science and Translational Research (CSTR) pilot program to RS. We thank the Texas A&M Institute for Genomic Medicine (TIGM) for providing MCU^−/−^ mice and Dr. Phillip A. West (TAMU College of Medicine) for useful discussions on mitochondrial biology. We thank Dr. Madesh Muniswamy (University of Texas Health Science Center at San Antonio) and Dr. Justin J. Wilson (Cornell University) for providing the MCU-specific inhibitor, Ru265.

