## Supplementary figures for "*In situ* characterization of calcium fluxes in astrocytic mitochondria from the mouse striatum and hippocampus"

**Abbreviated Title:** *In situ* characterization of astrocytic mitochondrial $Ca^{2+}$

**Authors:** Taylor E. Huntington[1,2] and Rahul Srinivasan[#1,2]

**Author addresses:** Department of Neuroscience & Experimental Therapeutics[1], Texas A&M University College of Medicine, 8447 Riverside Pkwy, Bryan, TX 77807-3260. Texas A&M Institute for Neuroscience (TAMIN) [2].

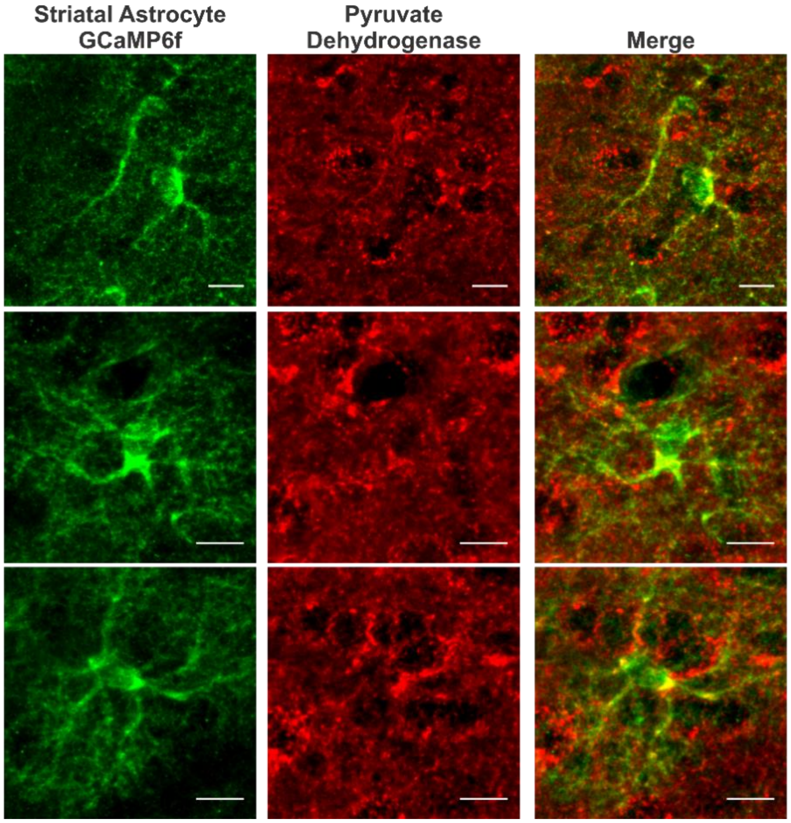


**Supplementary Figure 1. GfaABC$_1$D-mito-7-GCaMP6f co-localizes with pyruvate dehydrogenase.** Three representative confocal z-stack projections of mito-7-GCaMP6f expression in DLS astrocytic mitochondria co-stained with GFP (green) and pyruvate dehydrogenase (PDH) (red) antibodies. Merged images show clear co-localization of mito-7-GCaMP6f and PDH in DLS astrocytes. Scale bar = 10 µm.

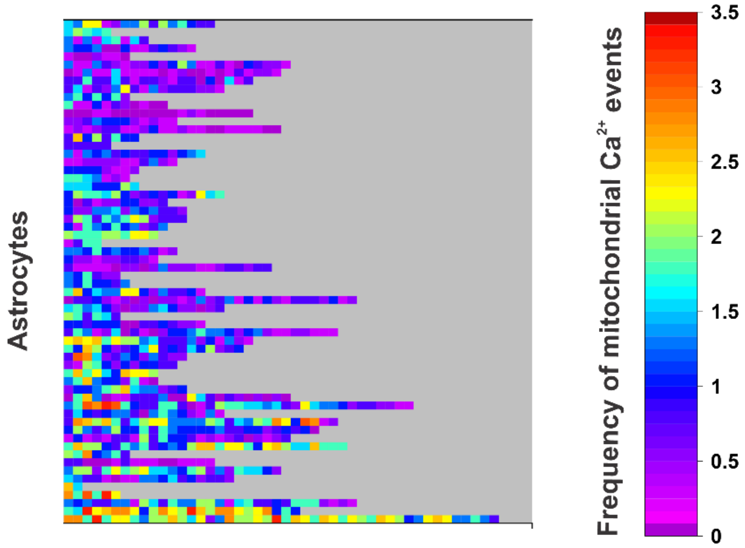


**Supplementary Figure 2.** Color coded distribution of mitochondrial $Ca^{2+}$ frequencies from 56 individual DLS astrocytes and 15 mice (n= 67 somatic, 336 branch, and 605 branchlet mitochondria). Each small colored box represents one mitochondrial ROI and each row is one astrocyte.

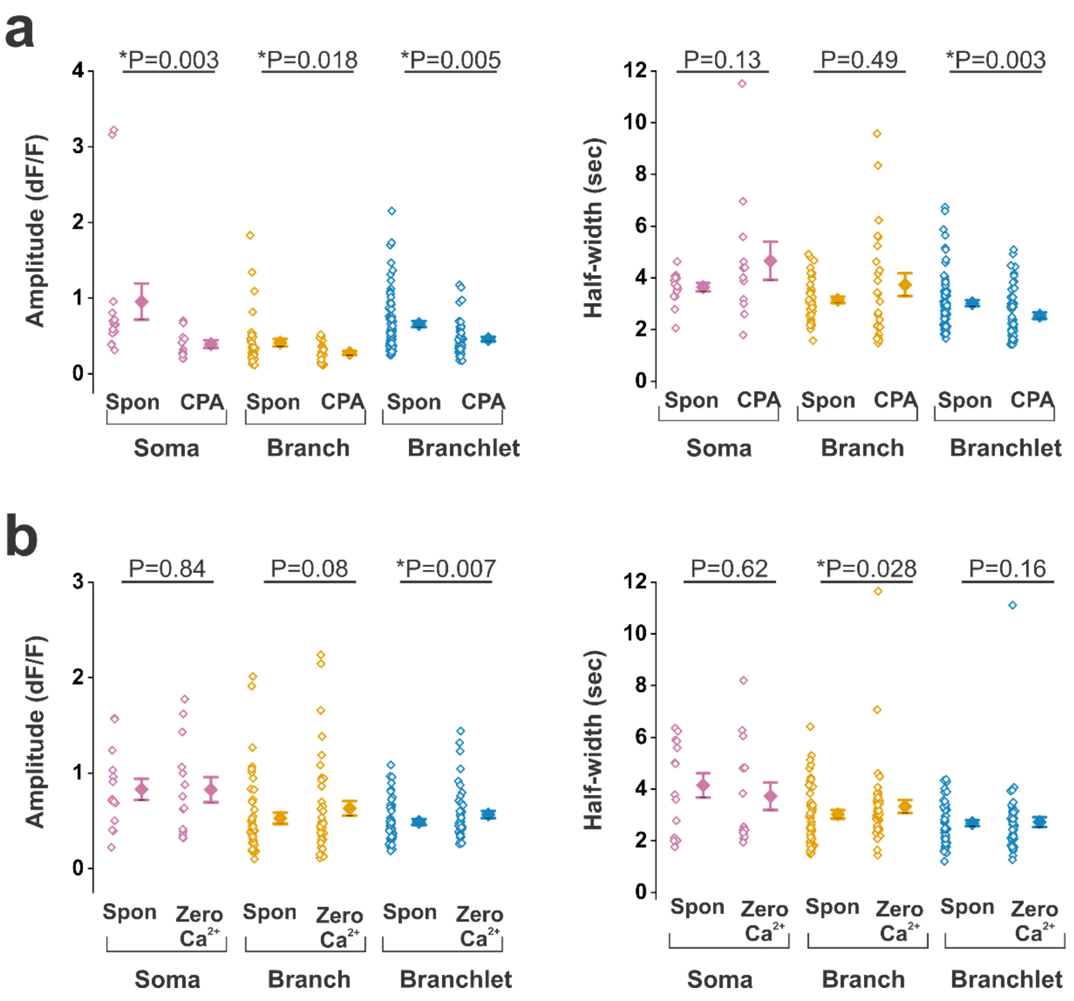


**Supplementary Figure 3. DLS astrocytic $Ca^{2+}$ event amplitude and half-width in response to CPA and Zero $Ca^{2+}$.** Population data and mean values of $Ca^{2+}$ event amplitude (left) and half-width (right) changes in DLS astrocytic mitochondria from somata (magenta), branches (orange), and branchlets (blue) with **a**, CPA from 7 astrocytes and 4 mice (n= 15 somatic, 43 branch, and 86 branchlet mitochondria) **b**, Zero $Ca^{2+}$ from 8 astrocytes and 4 mice (n= 12 somatic, 45 branch, and 59 branchlet mitochondria). Errors are ± s.e.m. For CPA data, p-values are based on Wilcoxon Signed Rank test. For Zero $Ca^{2+}$ the p-value for somatic mitochondrial amplitude is based on paired t-test, all other data sets are based on Wilcoxon Signed Rank test.

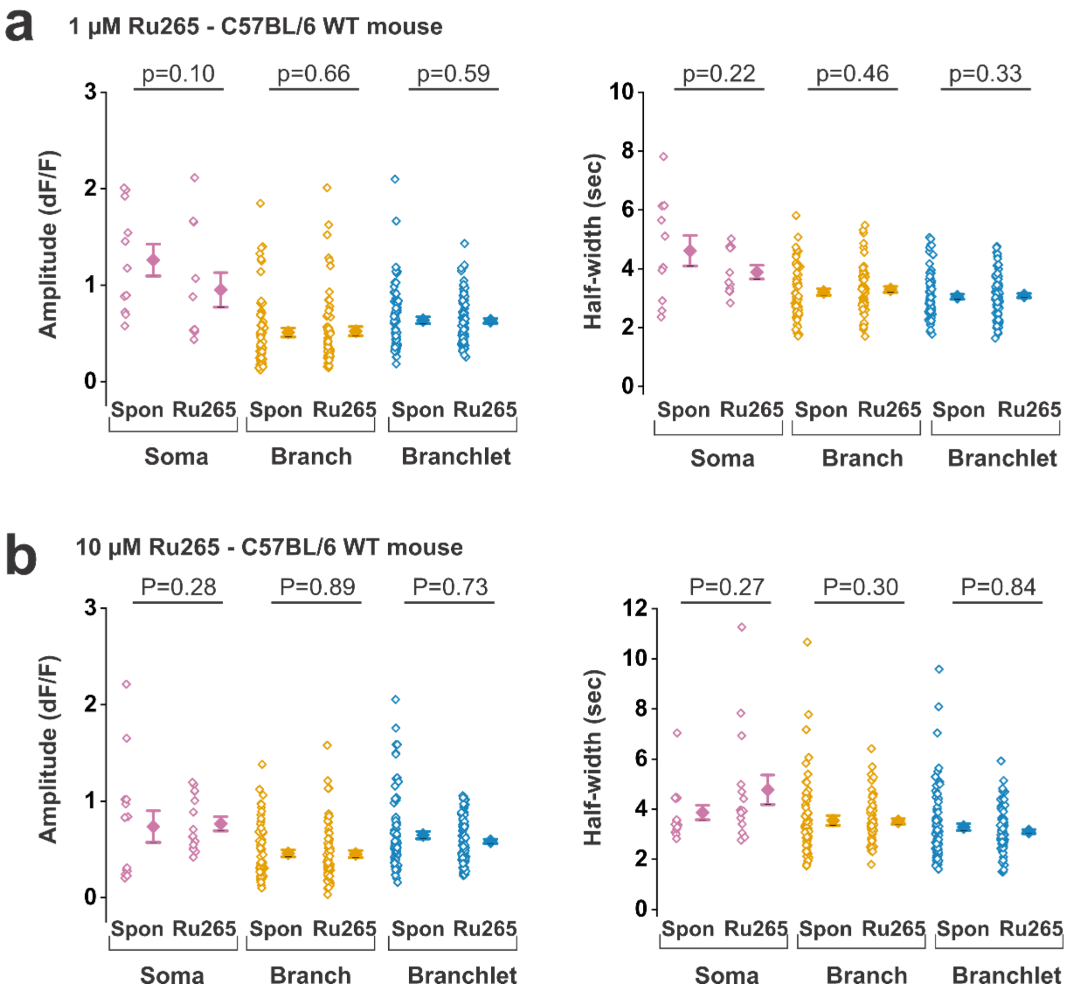


**Supplementary Figure 4. $Ca^{2+}$ event amplitude and half width responses to Ru265 in C57BL/6 WT mice. a**, Population data and mean values for DLS mitochondrial $Ca^{2+}$ event amplitude (left) and half-width (right) in response to 1 µM Ru265 from 7 astrocytes and 4 mice (n= 11 somatic, 62 branch, and 80 branchlet mitochondria). **b**, As in **a**, but in response to 10 µM Ru265 from 7 astrocytes and 4 mice (n= 13 somatic, 57 branch, and 90 branchlet mitochondria). Colors are somatic (magenta), branch (orange), or branchlet (blue) mitochondria. Errors are ± s.e.m. For 1 µM Ru265 data, the p-value for somatic mitochondria half-width is based on paired t-test, all other p-values are based on Wilcoxon Signed Rank test. For 10 µM, all p-values are based on Wilcoxon Signed Rank test.

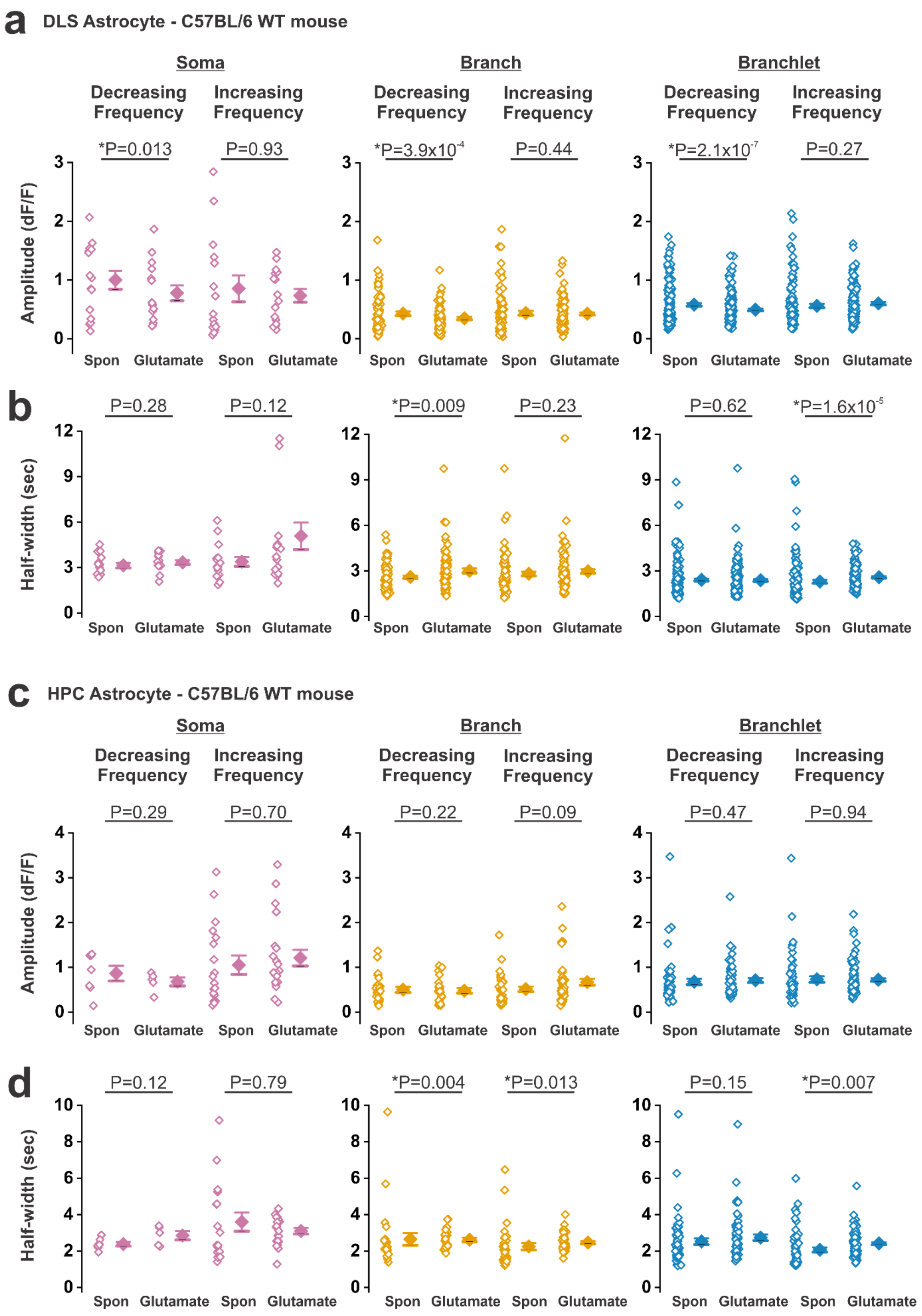


**Supplementary Figure 5. DLS and HPC mitochondrial $Ca^{2+}$ event amplitude and half-width changes in response to glutamate.** Population data and mean values for $Ca^{2+}$ event **a**, amplitude and **b**, half-width changes in DLS astrocytic mitochondria before and after glutamate.

Subpopulations are further separated by event frequency decrease or increase from 23 DLS astrocytes and 12 mice (n= 15 decreasing and 16 increasing somatic, 75 decreasing and 110 increasing branch, and 165 decreasing and 135 increasing branchlet mitochondria) **c-d**, As in **a-b**, but from mitochondria in 17 CA1 astrocytes and 8 mice (n= 7 decreasing and 21 increasing somatic, 25 decreasing and 44 increasing branch, and 59 decreasing and 97 increasing branchlet mitochondria). Colors are somatic (magenta), branch (orange), or branchlet (blue) mitochondria. Errors are ± s.e.m. For data from DLS astrocytic mitochondria, decreasing frequency somatic mitochondria amplitude and half-width p-values are based on paired t-test. All other p-values for DLS data are based on Wilcoxon Signed Rank test. For HPC data, decreasing frequency somatic mitochondria amplitude and half-width p-values are based on paired t-test. All other p-values for HPC data are based on Wilcoxon Signed Rank test.

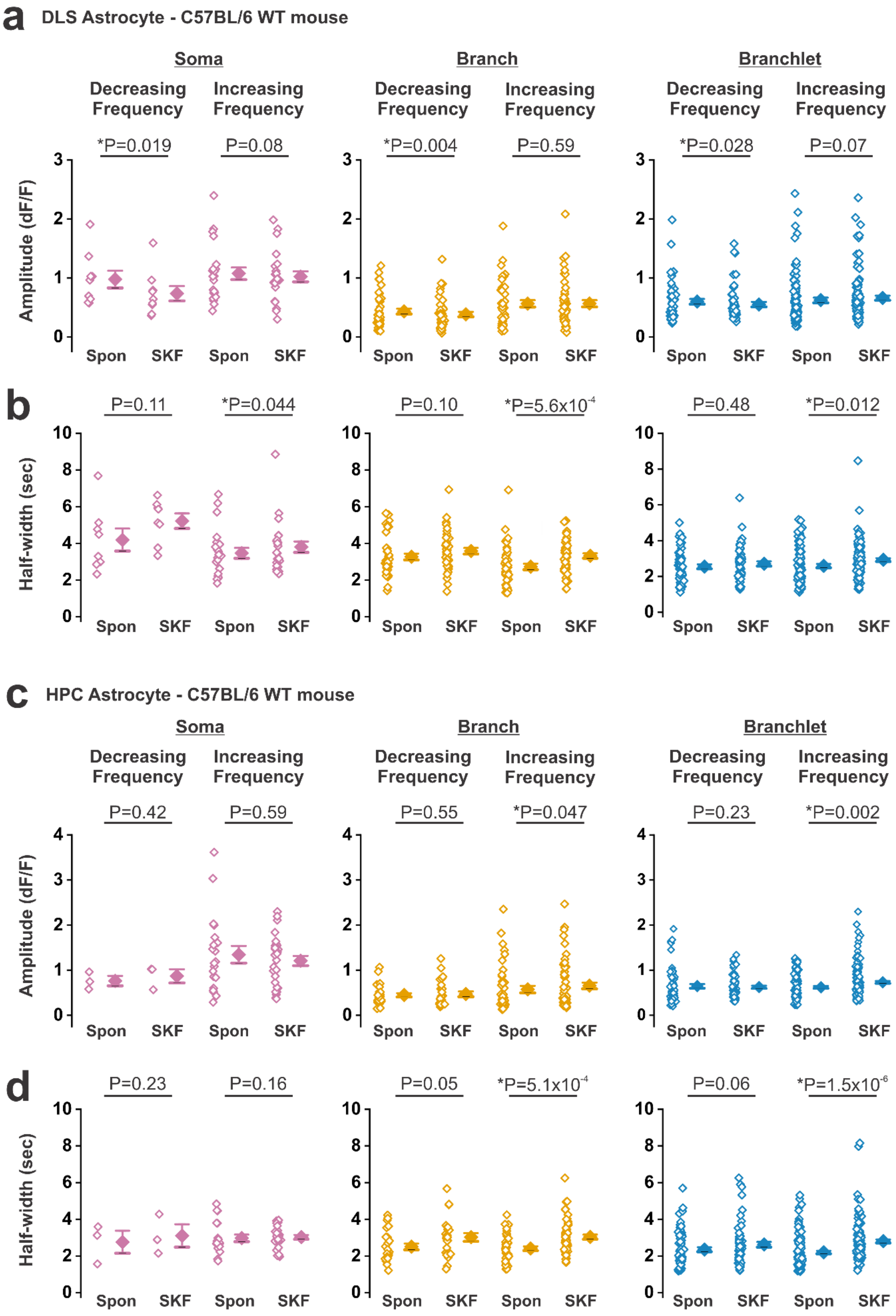


**Supplementary Figure 6. DLS and HPC mitochondrial $Ca^{2+}$ event amplitude and half-width changes in response to SKF-38393.** Population data and mean values for $Ca^{2+}$ event **a**, amplitude and **b**, half-width changes in DLS astrocytic mitochondria before and after SKF-

38393. Subpopulations are further separated by event frequency decreasing or increasing from 17 DLS astrocytes and 8 mice (n= 9 decreasing and 23 increasing somatic, 40 decreasing and 47 increasing branch, and 53 decreasing and 105 increasing branchlet mitochondria). **c-d**, As in **a-b**, but from mitochondria in 17 HPC CA1 astrocytes and 7 mice (n= 3 decreasing and 27 increasing somatic, 26 decreasing and 57 increasing branch, and 65 decreasing and 165 increasing branchlet mitochondria) responding to SKF-38393. Colors are somatic (magenta), branch (orange), or branchlet (blue) mitochondria. Errors are ± s.e.m. For data from DLS astrocytic mitochondria, decreasing frequency somatic mitochondria amplitude and half-width p-values are based on paired t-test. All other p-values for DLS data are based on Wilcoxon Signed Rank test. For HPC data, increasing frequency somatic mitochondria amplitude and decreasing frequency somatic mitochondria half-width p-values are based on paired t-test. All other data sets for HPC are based on Wilcoxon Signed Rank test.

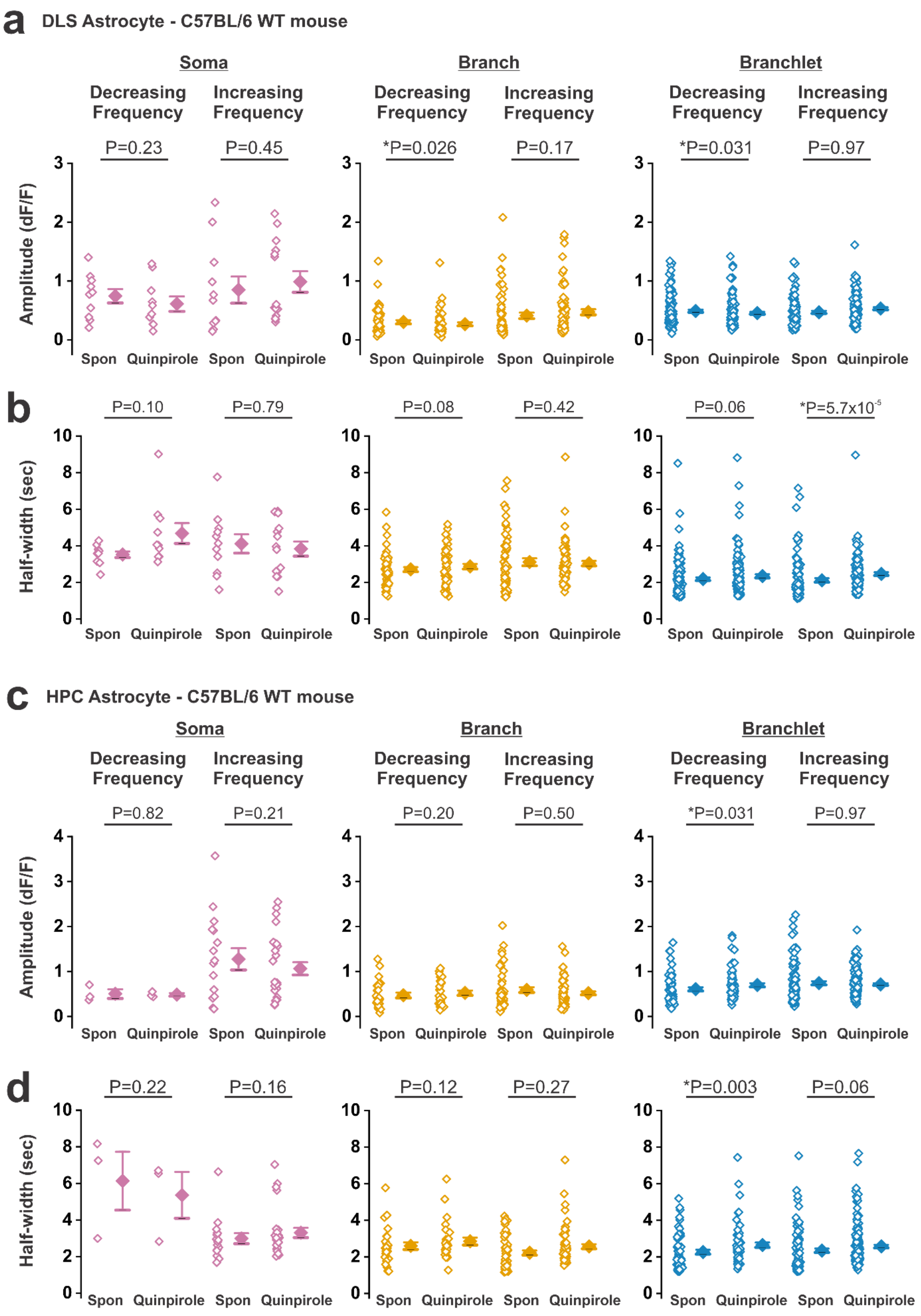


**Supplementary Figure 7. Ca$^{2+}$ event amplitude and half width responses to quinpirole in DLS and HPC astrocytic mitochondria.** Population data and mean values for Ca$^{2+}$ event **a**, amplitude and **b**, half-width changes in DLS astrocytic mitochondria before and after quinpirole.

Subpopulations are further separated by event frequency decreasing or increasing from 21 DLS astrocytes and 11 mice (n= 10 decreasing and 14 increasing somatic, 51 decreasing and 67 increasing branch, and 129 decreasing and 121 increasing branchlet mitochondria). **c-d**, As in **a-b**, but from mitochondria in 16 HPC CA1 astrocytes and 7 mice (n= 3 decreasing and 26 increasing somatic, 26 decreasing and 60 increasing branch, and 63 decreasing and 157 increasing branchlet mitochondria) responding to quinpirole. Colors represent somatic (magenta), branch (orange), or branchlet (blue) mitochondria. Errors are ± s.e.m. For DLS data, decreasing frequency somatic mitochondria amplitude and increasing frequency somatic mitochondria half-width p-values are based on paired t-test. All other DLS data sets are based on Wilcoxon Signed Rank test. For HPC data, decreasing frequency somatic mitochondria amplitude and half-width p-values are based on paired t-test. All other p-values for HPC data are based on Wilcoxon Signed Rank test.
