## Supplementary movie captions for "*In situ* characterization of calcium fluxes in astrocytic mitochondria from the mouse striatum and hippocampus"

**Abbreviated Title:** *In situ* characterization of astrocytic mitochondrial $Ca^{2+}$

**Authors:** Taylor E. Huntington[1,2] and Rahul Srinivasan[#1,2]

**Author addresses:** Department of Neuroscience & Experimental Therapeutics[1], Texas A&M University College of Medicine, 8447 Riverside Pkwy, Bryan, TX 77807-3260. Texas A&M Institute for Neuroscience (TAMIN) [2].

### Supplementary movie captions

**Supplementary movie 1**
DLS astrocyte from an adult WT C57BL/6 mouse with spontaneous $Ca^{2+}$ events in mitochondria. Frame rate is 0.8 seconds per frame.

**Supplementary movie 2**
DLS astrocyte from an adult WT C57BL/6 mouse with mitochondrial $Ca^{2+}$ events before and after exposure to CPA. Frame rate is 1 second per frame.

**Supplementary movie 3**
DLS astrocyte from an adult WT C57BL/6 mouse with mitochondrial $Ca^{2+}$ events before and after zero $Ca^{2+}$. Frame rate is 1 second per frame.

**Supplementary movie 4**
DLS astrocyte from an adult $MCU^{-/-}$ mouse with spontaneous mitochondrial $Ca^{2+}$ events. Frame rate is 0.8 seconds per frame.

**Supplementary movie 5**
DLS astrocyte from an adult WT C57BL/6 mouse with mitochondrial $Ca^{2+}$ events before and after administration of the MCU inhibitor, 10 µM Ru265. Frame rate is 1 second per frame.

**Supplementary movie 6**
HPC astrocyte with spontaneous mitochondrial $Ca^{2+}$ events from an adult WT C57BL/6 mouse. Frame rate is 0.8 seconds per frame.
